# All-viral tracing of monosynaptic inputs to single birthdate-defined neurons in the intact brain

**DOI:** 10.1101/2021.10.18.464781

**Authors:** R Irene Jacobsen, Rajeevkumar R Nair, Horst A Obenhaus, Flavio Donato, Torstein Slettmoen, May-Britt Moser, Edvard I Moser

## Abstract

Neuronal firing patterns are the result of inputs converging onto single cells. Identifying these inputs, anatomically and functionally, is essential to understand how neurons integrate information. Single-cell electroporation of helper genes and subsequent local injection of recombinant rabies viruses enable precise mapping of inputs to individual cells in superficial layers of the intact cortex. However, access to neurons in deeper structures requires more invasive procedures, including removal of overlying tissue. We have developed a method that through a combination of virus injections allows us to target ≤4 hippocampal cells 48% of the time and a single cell 16% of the time in wildtype mice without the use of electroporation or tissue aspiration. We identify local and distant monosynaptic inputs that can be functionally characterised *in vivo*. By expanding the toolbox for monosynaptic circuit tracing, this method will help further our understanding of neuronal integration at the level of single cells.

## Motivation

Identifying the inputs to a neuron is essential to understand its output. While this is possible with current methods in many neocortical areas, reaching cells in deeper brain regions *in vivo* requires removal of tissue that may be part of the circuit under investigation. We therefore developed a virus-based method that enables individual cells in sub-neocortical areas of wildtype mice to be targeted and their monosynaptic inputs identified without removing overlaying tissue.

## Introduction

The firing properties of individual neurons are the result of a convergence of multiple inputs. Genetically engineered rabies viruses enable the identification of monosynaptic inputs to populations of cells (Wickersham *et al*., 2007; Reardon *et al*., 2016; Ciabatti *et al*., 2017; Chatterjee *et al*., 2018) and have had a significant impact on our ability to dissect neural circuits. However, to understand the input/output transformation in neurons we need to map the inputs to individual cells, which is not trivial to achieve *in vivo*. Currently, it is possible to target a single such ‘starter cell’ in superficial cortical layers *in vivo* using single-cell electroporation (Marshel *et al*., 2010; Wertz *et al*., 2015; Rompani *et al*., 2017; Rossi, Harris and Carandini, 2020), patch clamping (Rancz *et al*., 2011; Vélez-Fort *et al*., 2014) or stamping (Schubert *et al*., 2017) techniques. However, targeting cells in deeper structures with these methods requires invasive procedures, including the removal of overlaying tissue (Rompani *et al*., 2017) that may be part of the circuit under investigation.

We therefore set out to develop a method that allows us to target individual neurons for monosynaptic rabies tracing in deep brain areas without the need for tissue aspiration. To demonstrate our method, we chose the hippocampal tri-synaptic circuit as this is a network that has received considerable attention due to its importance for learning, memory, and navigation. As a result, the different hippocampal subfields and firing properties of the principal cell types within it, as well as the gross connectivity of the circuit, are well understood, yet the constellation of inputs an individual cell receives remains unknown. This is in part because rabies-mediated input mapping of single hippocampal cells requires the removal of substantial amounts of overlaying tissue, including parts of CA1 to reach the dentate gyrus and CA3, which disrupts the tri-synaptic circuit. To target single cells and identify their monosynaptic inputs in the intact hippocampus, we first performed an *in utero* injection of a virus carrying Cre recombinase to achieve sparse expression of Cre in the adult brain. Subsequently, by locally injecting a custom helper virus that reaches only a single neuron within this sparsely Cre-labelled population, we can, in a single cell, express both the receptor and G-protein necessary for rabies infection and spread, respectively. We show that our approach enables the identification of both local and distant inputs to these individual starter cells within the hippocampal-entorhinal circuit. Further, our *in utero* approach enables cells to be targeted based on their birthdate during development, making it possible to investigate inputs to cells with specific developmental origins. We demonstrate how this method can be used to determine the functional properties of monosynaptic inputs to such cell populations with *in vivo* electrophysiology combined with optogenetic tagging, or calcium imaging. Specifically, we provide proof of concept for determining the spatial tuning properties of cells in the medial entorhinal cortex (MEC) that project directly to birthdate-defined cell populations in the hippocampus.

By enabling individual cells in sub-neocortical brain structures to be targeted for monosynaptic input mapping in a minimally invasive manner, this method extends rabies virus-based circuit-mapping tools to deeper regions. This will in turn expand our understanding of afferent connectivity and the functional integration of inputs that occurs in single cells in these circuits.

## Results

### A virus-based method for targeting a single cell and identifying its inputs

We first designed and produced the viruses necessary to perform rabies-mediated tracing based on the CVS-N2c strain, as this strain spreads to more input cells and is less cytotoxic than the SAD-B19 strain (Reardon *et al*., 2016). We produced three different EnvA-pseudotyped, G-protein deleted CVS-N2c recombinant rabies viruses expressing a gene of interest (RABV-GOI) each. Specifically, tdTomato (RABV-tdTomato), channelrhodopsin 2 (RABV-ChR2-YFP) or GCaMP6f (RABV-GCaMP6f), to enable anatomical circuit mapping, optogenetic tagging, and calcium imaging of input cells, respectively (**Figure 1A**). These pseudotyped rabies viruses require the TVA receptor to enter cells and a G-protein to spread to input cells (**Figure S1A**). We incorporated both TVA and the G-protein from the CVS-N2c strain into a single Cre-dependent adeno associated virus (‘helper virus’, AAV-hS-FLEX-TVA-HA-N2cG, **Figure 1A**), thereby preventing unspecific and undetected starter cell labelling that can occur if these components are introduced separately using multiple helper viruses, for example by resulting in secondary jumping from cells that express G but not TVA. We selected an HA-tag as a surrogate marker instead of a fluorescent one to facilitate efficient packaging of viral particles. Further, the transgenes were separated by 2A self-cleaving peptide sequences (de Felipe *et al*., 2006; Tang *et al*., 2009), providing 2A peptide epitopes for immunodetection. These strategies enabled us to perform immunohistological identification of HA or 2A ‘helper tags’ (**Figure 1B**) in cells infected with our new, single helper virus and ensured compatibility with rabies viruses expressing different fluorescent payloads.

**Figure 1:**
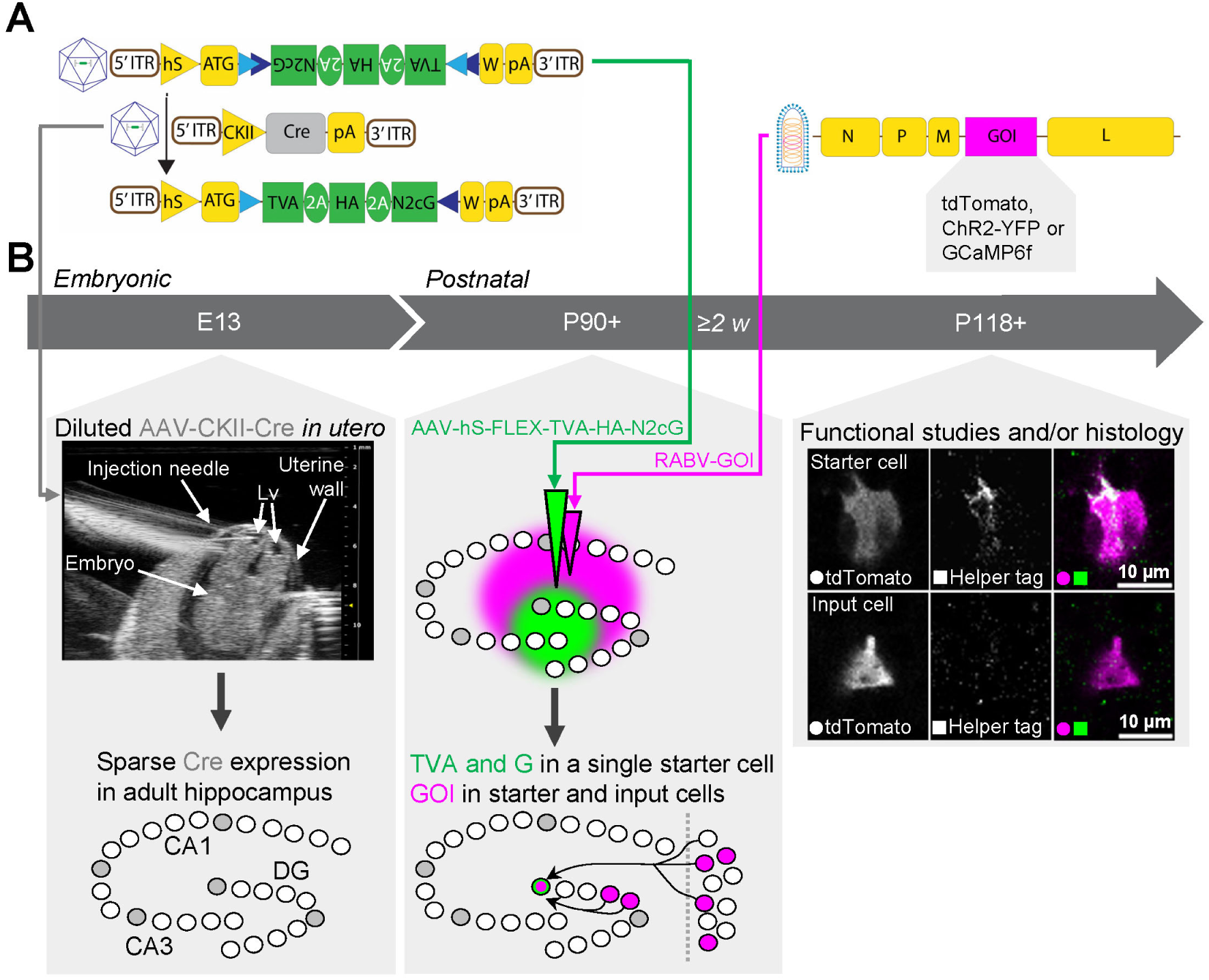
A virus-based method for targeting a single cell and identifying its inputs. **A)** Schematics of the viral vectors used in this project: AAV-hS-FLEX-TVA-HA-N2cG (top left), AAV-CKII-Cre (middle left), the top left vector after Cre-mediated recombination (bottom left) and EnvA-pseudotyped ΔG-CVS-N2c rabies virus (RABV-GOI, right, EnvA: envelope protein from the subgroup-A avian sarcoma and leukosis virus). ITR: Inverted Terminal Repeat, hS: human synapsin promoter, ATG: start codon, blue overlapping triangles: FLEX cassette consisting of loxP and lox2272 sites, N2cG: CVS-N2c rabies glycoprotein, 2A: self-cleaving peptide element, HA: hemagglutinin tag, TVA: start codon removed avian-specific receptor, W: woodchuck hepatitis virus post-transcriptional regulatory element, pA: bovine growth hormone polyadenylation signal, CKII: CaM kinase II promoter, N, P, M, L: genes coding the necessary viral proteins for the rabies virus, GOI: gene of interest. **B)** Timeline of experimental protocol that starts when a male and female mouse are introduced for breeding at E0 (embryonic day 0) for up to 24 hours. E13: a diluted AAV carrying Cre is injected into one of the lateral ventricles (Lv) of a mouse embryo under ultrasound guidance, resulting in Cre-expression (grey cells in schematic) in a sparse number of cells in the adult hippocampus. P90+: in combination with sparse Cre expression, a small volume injection of a single rabies helper virus (green) into the hippocampus leads to the expression of TVA (the receptor for EnvA-pseudotyped rabies virus) and G (N2c rabies glycoprotein that is necessary for spread of the rabies virus) in a single hippocampal cell. A large volume injection of a G-protein deleted, EnvA-pseudotyped rabies virus (magenta) in the same area weeks later (≥2 weeks) results in infection of the single TVA and G expressing cell (green and magenta cell in schematic) and spread of the rabies virus to monosynaptically connected input cells (magenta-only cells) both locally within the hippocampus and in distant regions, such as the MEC. P118+: starter and input cells can be distinguished with antibody staining against HA or 2A epitope proteins introduced by the helper virus (i.e. providing a ‘helper tag’). Cells that show colocalization of both the helper tag and the GOI introduced by the rabies virus (such as tdTomato) are identified as starter cells, while cells only expressing the GOI are identified as input cells. Images are single z-plane confocal images. E: embryonic day, P: postnatal day.

Next, we used these viruses to label the monosynaptic inputs to cells in the hippocampus by first expressing Cre in wildtype mice. To target a single cell, we performed *in utero* injections of an AAV carrying Cre under the control of the CaMKII promoter (AAV-CKII-Cre) diluted in PBS at embryonic day 13 (E13, **Figure 1B, Table S1**), when cells in the hippocampal subfields are born (Angevine, 1965; Deguchi *et al*., 2011) and the lateral ventricles are still clearly visible (**Figure 1B**). Using ultrasound to guide the pipette, we targeted one of the lateral ventricles and injected approximately 300 nl in every embryo. Due to the rapid turnover of cerebrospinal fluid and hence the restricted temporal exposure of cells lining the ventricles to the virus (Donato *et al*., 2017), only a sparse number of cells express Cre in the adult hippocampus of these animals (**Figure S1B**). Next, we injected approximately 50 nl of the helper virus carrying Cre-dependent TVA and G under the control of the human synapsin promoter (AAV-hS-FLEX-TVA-HA-N2cG) unilaterally into the adult hippocampus, targeting the CA3 and dentate gyrus (**Figure 1B**). At least two weeks later, a separate injection of the pseudotyped rabies virus carrying tdTomato (RABV-tdTomato) resulted in infection of TVA expressing cells (‘starter cells’) and, with time, G-protein-mediated spread of the rabies virus carrying tdTomato to monosynaptically connected input cells (**Figure 1B**). To determine the number of starter cells targeted, we perfused all animals within one month of the rabies injection (if we waited longer the infected cells started to look unhealthy, **Figure 7A**). We next sectioned the ipsilateral hemisphere, keeping all sections and taking care to maintain their original order. Antibodies were then used to label cells expressing a helper tag (**Figure 1B**) as well as those expressing tdTomato (to ensure that rabies infected cells were not missed due to weak tdTomato expression). All tdTomato expressing cells within the dorsal hippocampus were scanned at 40x with a confocal microscope to check for co-expression with the helper tag. Cells labelled with both the helper tag and tdTomato were identified as starter cells while cells only expressing tdTomato were classified as input cells (**Figure 1B**). Due to the difficulties associated with antibody detection of helper tags (see Discussion), we split the animals into two groups (see STAR Methods): those in which starter cells could be clearly detected (‘conclusive’ animals) and those in which it was ambiguous if starter cells were present (‘inconclusive’ animals).

With this approach, we could unambiguously target four or fewer starter cells in 48% (15/31) of animals (**Figure 2A**). Within this group, a single starter cell was labelled 33% (5/15) of the time (**Figure 2A**), resulting in an overall success rate of 16% (5/31) for targeting a single cell in the hippocampus. As expected, longer expression times of the rabies virus increased the number of rabies positive cells (**Figure 2B**), the anatomical range over which they were found in the dorsal hippocampus (**Figure S2A**), as well as the number of input cells labelled in the more distant MEC (**Figure S2B**). Hence, we can identify both local hippocampal and distant MEC inputs to single cells in the hippocampus (**Figure 3 and 4**). In addition, the identity of the inputs reflected the identity of the starter cell (**Figure 3 and 4**), which is an important indication that the rabies virus is indeed transsynaptic. For example, the constellation of inputs that a single pyramidal neuron and a single interneuron received was very different despite them being located in neighbouring regions (distal CA3 and CA2) in the hippocampus (**Figure 3**).

**Figure 2:**
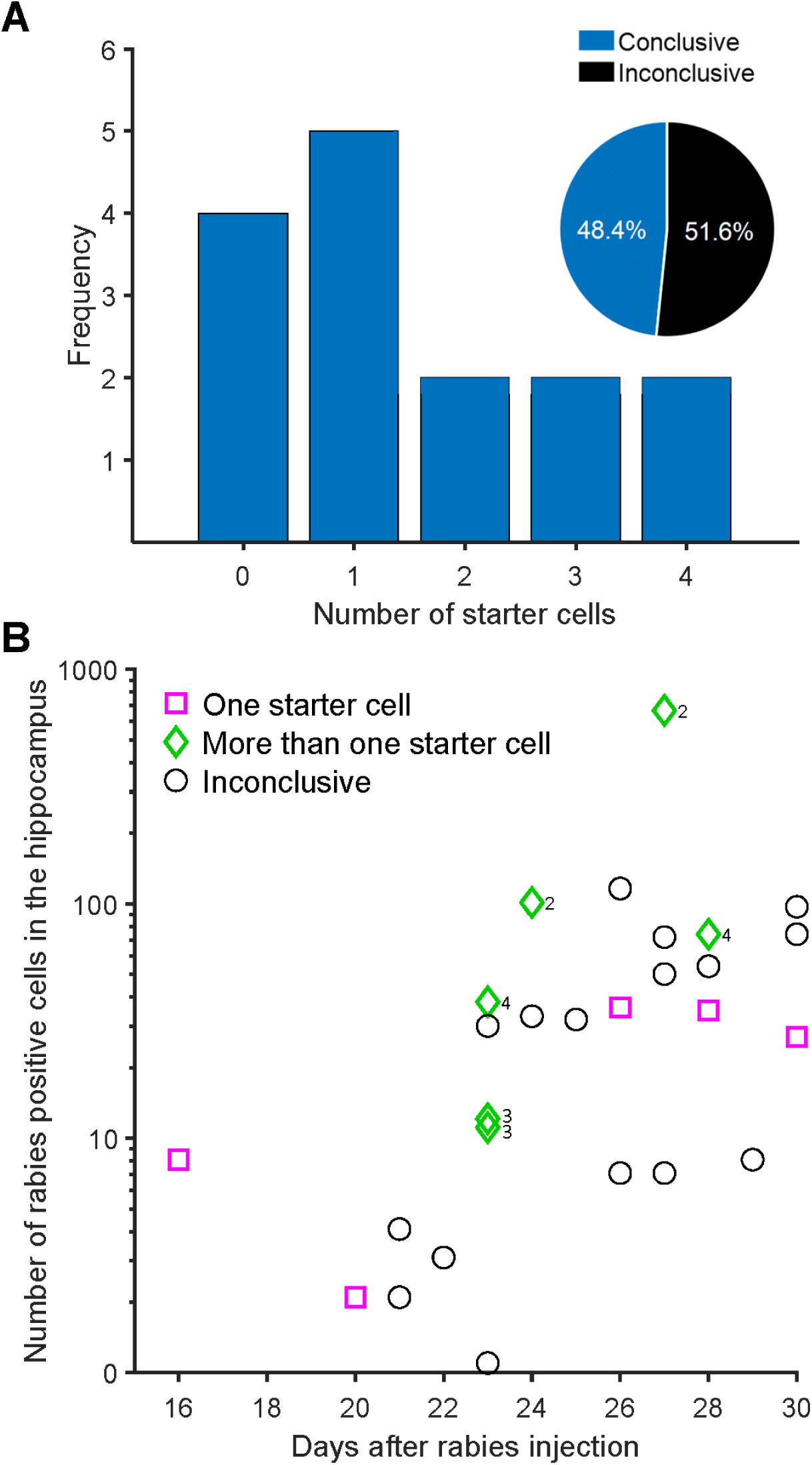
Success rate and number of rabies-positive cells in the dorsal hippocampus. **A)** The frequency of conclusive starter cells observed with our method. The pie chart inset indicates the proportion of animals in which starter cells could be conclusively identified based on unambiguous staining of helper virus proteins (n = 31 animals, 15 of which were ‘Conclusive’). **B)** The total number of rabies-positive cells observed in the ipsilateral dorsal hippocampus increases as a function of days after the rabies injection (R = 0.58, p = 0.0014, Spearman’s rho, n = 27 animals). Numbers next to diamonds indicate the number of starter cells in the ‘More than one starter cell’ category. Note: the y-axis follows a log scale and animals with zero starter cells are not included as rabies-positive cells are absent. See also Figure S2.

**Figure 3:**
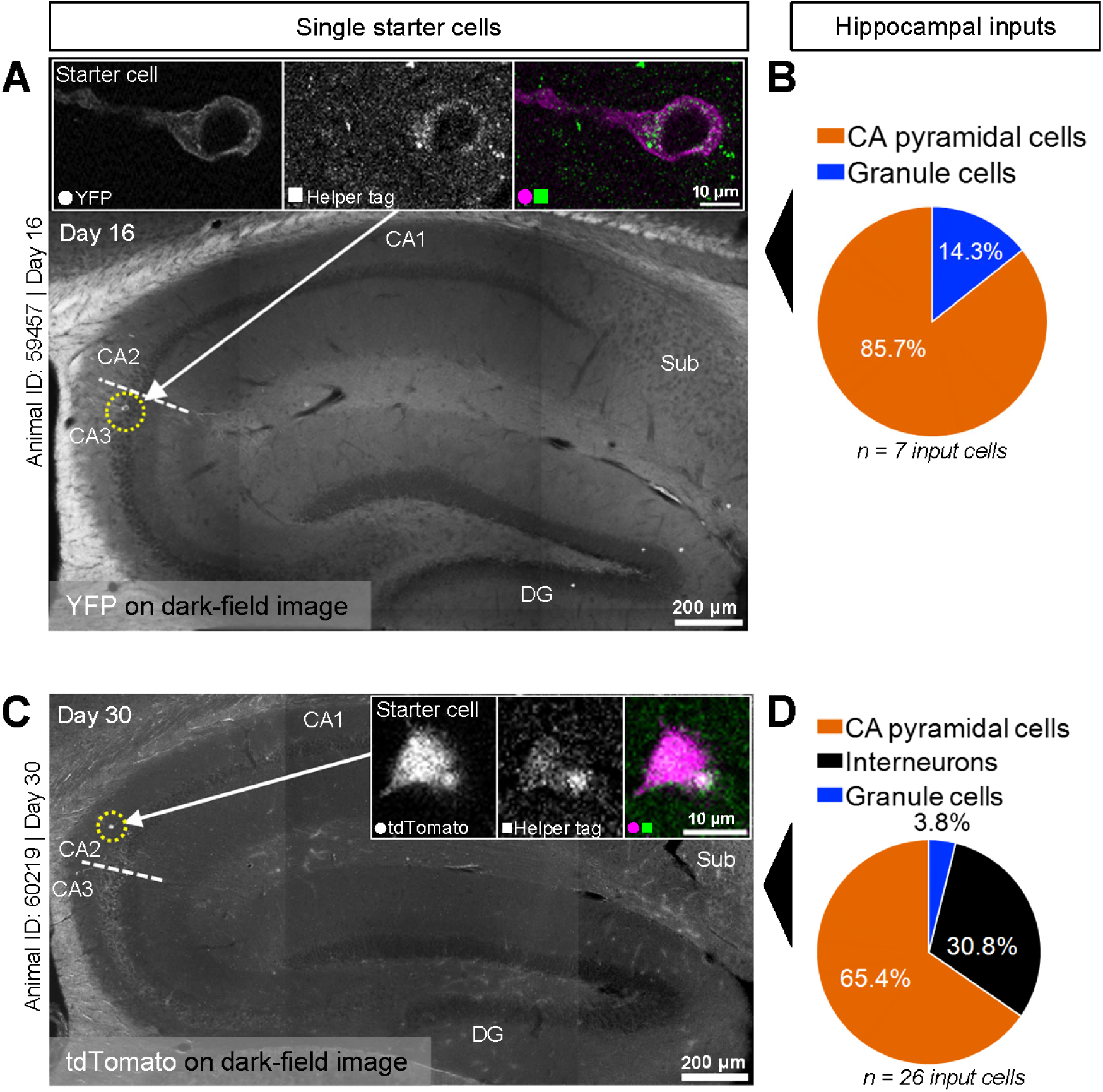
Distribution of input cells reflects identity of single starter cell. Examples of single starter cells are shown in the left column and their inputs in the right column. **A)** Examples of single starter cells are shown in the left column and their inputs in the right column. Dark-field image of the dorsal hippocampus with epifluorescence of a single starter cell overlaid and highlighted with an arrow and a yellow dotted circle. The cell was identified as a CA3 pyramidal cell based on its location within the pyramidal cell layer, cell morphology, and the presence of mossy fibres. The white dotted line indicates the border between CA3 and CA2 as defined by the presence and absence of mossy fibres, respectively. Image insets are single z-plane confocal images showing colocalization of antibody staining for a tag introduced by the helper virus and YFP introduced by the rabies virus (RABV-ChR2-YFP). **B)** Pie chart showing the proportions of hippocampal input cell types to the single starter cell in A) 16 days after the rabies injection (n = 7 input cells to a single starter cell). **C)** Dark-field image of the dorsal hippocampus with epifluorescence of a single starter cell in the stratum oriens overlaid and highlighted with an arrow and a yellow dotted circle. Based on its location and cell morphology, this was identified as an inhibitory neuron. The white dotted line indicates the border between CA3 and CA2 as defined by the presence and absence of mossy fibres, respectively. Image insets are single z-plane confocal images showing colocalization of antibody staining for a tag introduced by the helper virus and tdTomato introduced by the rabies virus (RABV-tdTomato). **D)** Pie chart showing the proportions of hippocampal input cell types to the single starter cell in C) 30 days after the rabies injection (n = 26 input cells to a single starter cell). DG: dentate gyrus, Sub: subiculum.

**Figure 4:**
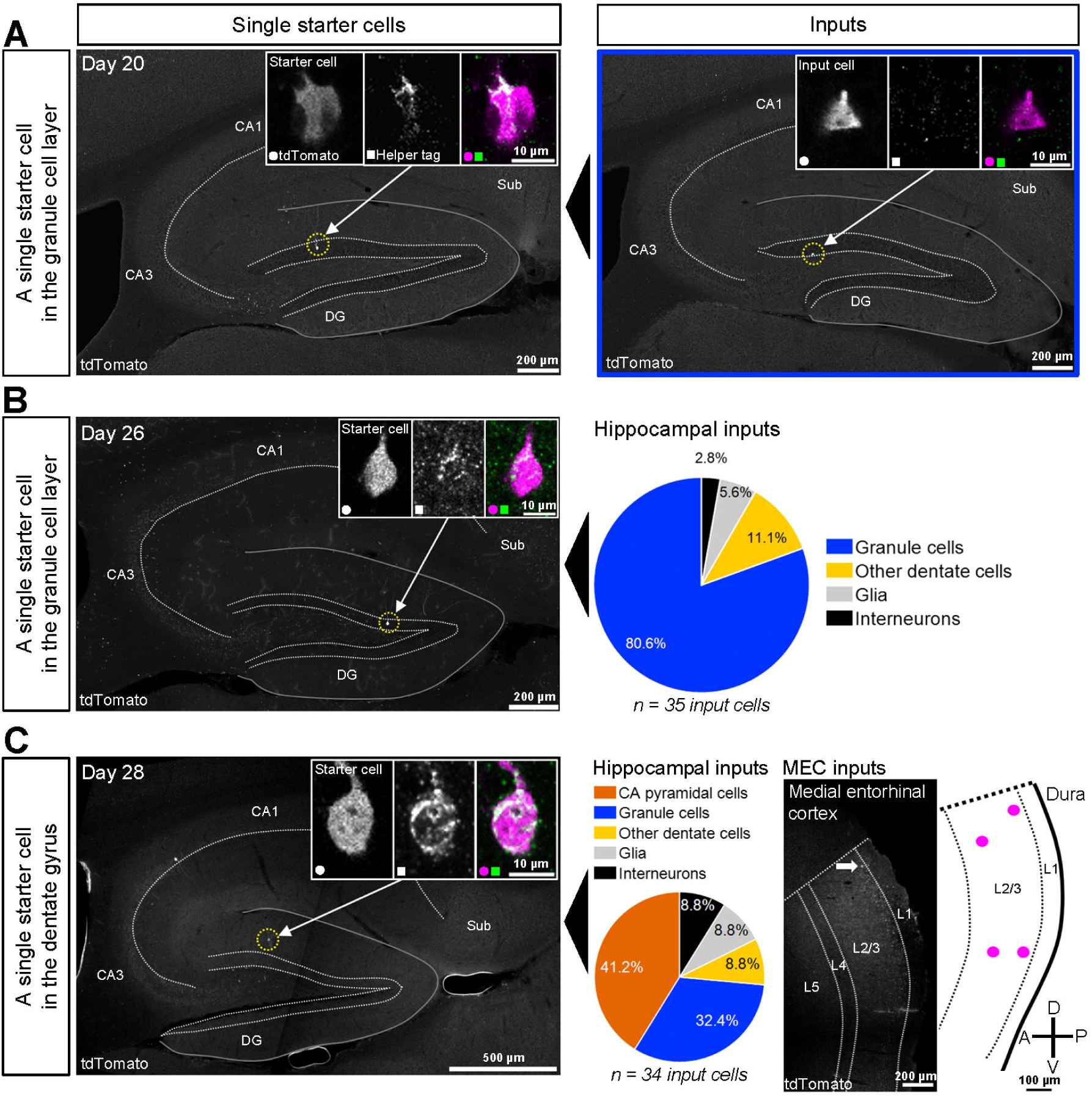
Local and distant inputs to three different single starter cells in the dentate gyrus. **A)** Confocal maximum intensity projection showing a single starter cell in the dentate gyrus granule cell layer in the dorsal hippocampus highlighted with an arrow and a yellow dotted circle (left). 20 days after the rabies injection, a single hippocampal input cell (right, also a granule cell and highlighted with an arrow) is found >320 µm away, along the longitudinal axis, from the starter cell. Image insets are single z-plane confocal images showing colocalization of antibody staining for a tag introduced by the helper virus and tdTomato introduced by the rabies virus (left insets; starter cell) and only tdTomato expression (right insets; input cell, n = 1 input cell to a single starter cell). **B)** Confocal maximum intensity projection showing a single starter cell in the dentate gyrus granule cell layer in the dorsal hippocampus highlighted with an arrow and a yellow dotted circle (left). Image insets are single z-plane confocal images showing colocalization of antibody staining for a tag introduced by the helper virus and tdTomato introduced by the helper virus (starter cell). The pie chart (right) shows the proportions of hippocampal input cell types to the cell highlighted on the left 26 days after the rabies injection (n = 35 input cells to a single starter cell). **C)** Epifluorescence images showing a single starter cell in the dentate gyrus of the dorsal hippocampus highlighted with an arrow and a yellow dotted circle (left, potentially a molecular layer perforant path associated interneuron) and one of its input cells in the dorsal MEC (middle right, arrow). Image insets are single z-plane confocal images showing colocalization of antibody staining for a tag introduced by the helper virus and tdTomato introduced by the rabies virus (left, starter cell). The pie chart (middle left) shows the proportions of hippocampal input cell types to the single starter cell highlighted on the left 28 days after the rabies injection (n = 34 input cells in the hippocampus to a single starter cell). A schematic (right) illustrates the location of all input cells to the same single starter cell across 400 µm of the medio-lateral extent of the MEC (n = 4 input cells in the MEC to a single starter cell in the hippocampus). A: anterior, D: dorsal, P: posterior, V: ventral. DG: dentate gyrus, Sub: subiculum.

### Targeting single starter cells in the hippocampus highlights granule-granule cell connectivity

A striking finding from the animals in which we achieved single starter cell labelling of hippocampal granule cells, was the high proportion of inputs from other granule cells. After 20 days of rabies expression, one granule cell was found to receive input from another granule cell located several hundred micrometres away along the longitudinal axis of the hippocampus (**Figure 4A**). After 26 days (in a different animal) the virus had spread to a total of 35 input cells in the hippocampus, 81% of which were other granule cells (**Figure 4B**). While the starter cells themselves were located in the middle of the granule cell layer, indicating that they are mature granule cells, a substantial number of their inputs were found in the subgranular zone (1/1 granule cell inputs after 20 days, 11/28 after 26 days), which suggests that these are immature granule cells. Although this type of granule-granule cell connection was unexpected, weak connections from adult-born to mature granule cells have been described (Drew *et al*., 2016) and connections going in the opposite direction have previously been identified with rabies tracing (Vivar *et al*., 2012). Additional studies are necessary to confirm our preliminary findings, but in combination with the mentioned literature they may suggest that in the case of hippocampal granule cells, rabies viruses might preferentially target synapses shared with other granule cells.

### The functional properties of input cells to birthdate-defined cell populations can be determined with optogenetics *in vivo*

Our *in utero* approach provides unique opportunities for developmental studies. By varying the timepoint at which the virus carrying Cre is injected, different populations of birthdate-matched cells can be targeted in wildtype animals (Donato *et al*., 2017). To determine the functional properties of inputs to such cell populations, our protocol can be adapted to 1) increase the number of starter cells, and 2) include techniques for measuring neural activity.

The number of starter cells can be adjusted by varying the amount of the Cre-carrying virus injected *in utero* and the amount of helper virus injected in the adult brain. By increasing both, we were able to target populations of hippocampal cells born on different embryonic days for rabies virus-mediated tracing (E12: **Figure 5AB, S1A**; E14: **Figure 6A**). Inputs to these birthdate-defined populations can also be compared to the inputs that a general population of hippocampal cells receives, by leaving out the *in utero* injection and introducing the Cre-carrying virus along with the helper virus in the adult brain instead (**Figure S3A**). Both approaches led to broad labelling of these populations’ input cells in superficial MEC (**Figure 5B, 6A, S1A, S3A**), which we next sought to characterise functionally. To do so, we first tested the feasibility of using *in vivo* electrophysiology combined with optogenetic tagging. We injected a rabies virus carrying ChR2 (RABV-ChR2-YFP) in the dorsal hippocampus and implanted tetrodes with an optic fibre attached into the ipsilateral MEC (**Figure 5A**), a major input region containing cell types that have been well-characterised with *in vivo* electrophysiology (Rowland *et al*., 2016). In these experiments, input cells could be detected in the MEC from approximately two weeks after the rabies injection. We started each session by recording single unit activity in the MEC while the mice explored an open field environment. We found several functional cell types, including grid, head direction and border cells (**Figure 5C, S4**), indicating that the network as a whole was not affected by the widespread rabies expression in the entorhinal-hippocampal network. Next, we placed the mice in a separate holding box and shone a 473 nm laser through the implanted optic fibre to determine if any of these cells responded to laser stimulation, i.e., if they express ChR2 introduced by the rabies virus and therefore project directly to the hippocampus. Cells that responded robustly to laser stimulation with a 2-4 millisecond latency (demonstrating direct activation as opposed to later, secondary activation via stimulation of upstream inputs) included grid cells (**Figure 5C**). This finding confirms a previous study (Zhang *et al*., 2013) and indicates that rabies expression does not preclude the functional identification of input cells. The firing properties of an individual input cell were also found to remain stable for at least three days of continuous rabies expression (**Figure S3**) and responsive cells with clear spatial firing patterns could be identified for at least 19 days after the rabies injection (**Figure 5C**). Combined, these results show that it is possible to functionally identify and characterise distant inputs to developmentally defined cell populations through rabies-mediated optogenetic tagging *in vivo*.

**Figure 5:**
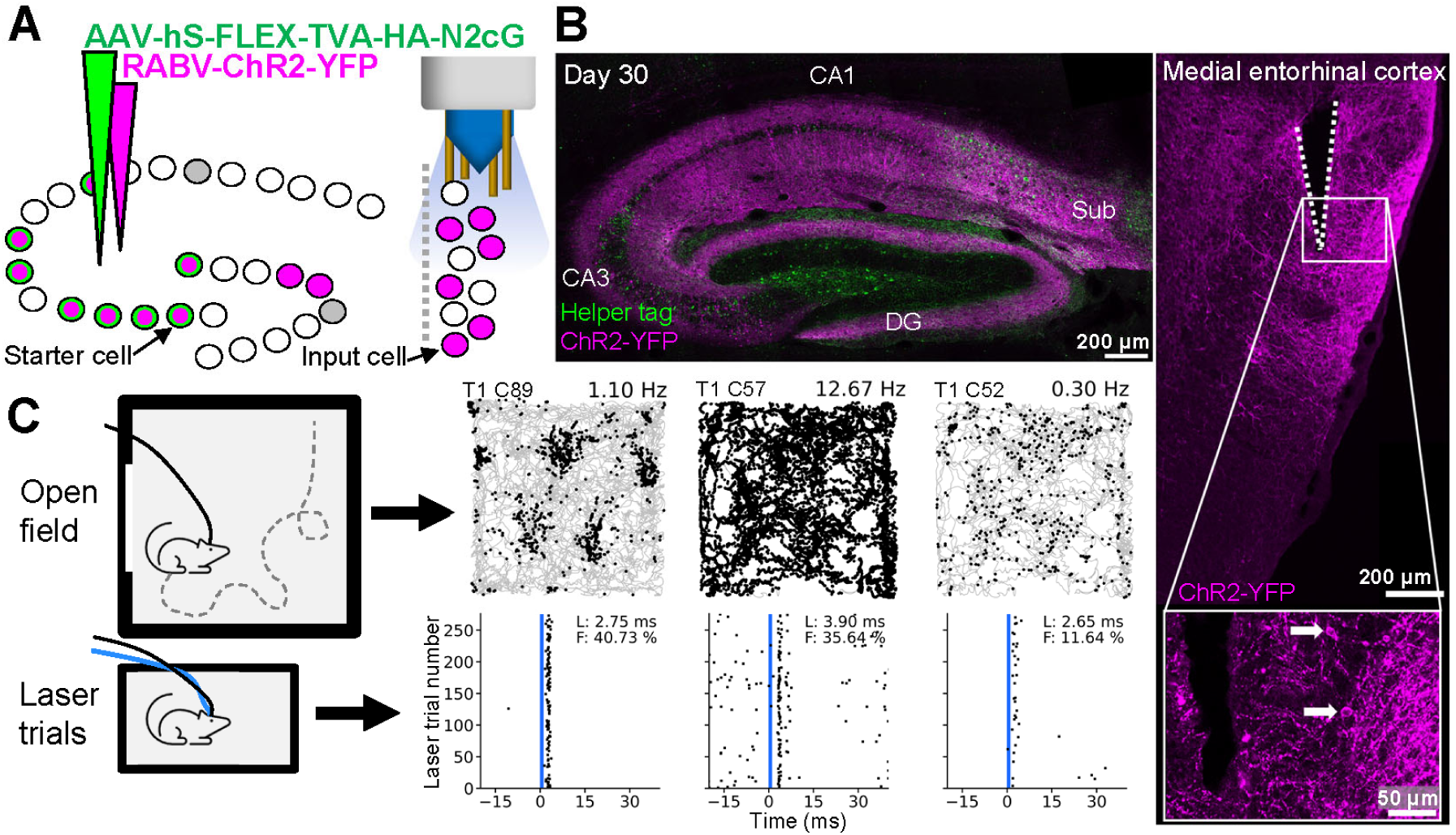
The functional properties of input cells to birthdate-defined cell populations can be determined with optogenetics in vivo. **A)** Schematic of experimental design. A population of birthdate-matched (E12) starter cells in the hippocampus was targeted by introducing an undiluted Cre virus during embryonic development and injecting a large volume (eight times higher than for the ‘single cell experiments’) of the helper virus in the adult hippocampus. In a separate surgery, a rabies virus carrying ChR2 and YFP was injected into the hippocampus and tetrodes with an optic fibre attached implanted in the ipsilateral MEC. **B)** Left: confocal maximum intensity projection of the dorsal hippocampus showing cells expressing helper proteins and/or ChR2-YFP 30 days after the rabies injection. Right: confocal maximum intensity projection showing part of the tetrode track (white dotted lines) and input cells expressing ChR2-YFP in the MEC. The inset is a single z-plane confocal image highlighting two example cells (arrows) near the tetrode track. DG: dentate gyrus, Sub: subiculum. **C)** The functional identity of cells was determined in an open field with a rectangular cue card on one wall while the mouse foraged for cookie crumbs. Afterwards, the mouse was placed in a separate holding box for laser stimulation trials to identify cells that express ChR2, which means they provide monosynaptic input to the hippocampus. Corresponding path and laser stimulation plots from three example cells that show robust and reliable responses to laser stimulation (blue vertical line) are shown. These include a grid cell (left, 19 days after rabies injection), a high firing rate cell (middle, 17 days after rabies injection) and a low firing rate cell (right, 17 days after rabies injection). Grey lines indicate the path of the mouse in the open field while black dots indicate locations (path plots) or timing relative to the laser stimulation (laser stimulation plots) at which units from an individual cell was recorded. Right-centred values above path plots show the average firing rate for the open field session. T: tetrode number, C: cell ID, L: latency, F: fidelity. See also Figure S3 and S4.

**Figure 6:**
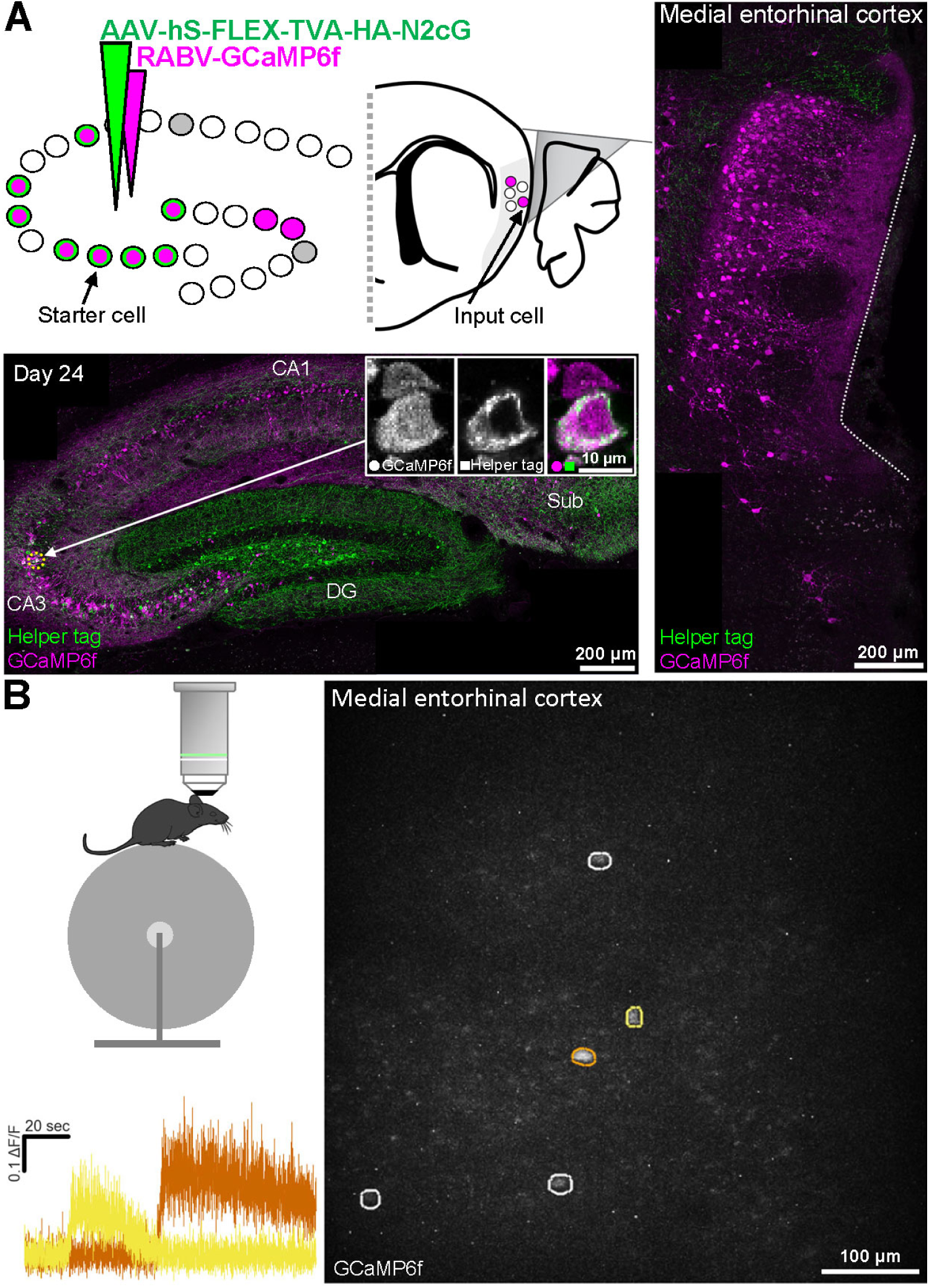
In vivo two-photon calcium imaging enables functional detection of distant input cells to birthdate-defined cell populations. **A)** Schematic of experimental design and resulting histology 24 days after the rabies injection. Top left: a population of birthdate-matched (E14) cells in the hippocampus was targeted to express TVA and G by injecting an undiluted virus carrying Cre in utero and a large volume of the rabies helper virus carrying TVA and G in the adult hippocampus. A rabies virus carrying GCaMP6f was used to label cells providing monosynaptic input to this cell population. These inputs were then visualised via a prism implanted between the MEC (brain region marked in grey) and the cerebellum. Bottom left: confocal maximum intensity projection of the dorsal hippocampus showing cells expressing helper proteins and/or GCaMP6f. Image insets are single z-plane confocal images showing colocalization of antibody staining for a tag introduced by the helper virus and GCaMP6f introduced by the rabies virus in one example cell (i.e. a starter cell) and only GCaMP6f expression in a neighbouring cell (i.e. an input cell) highlighted with an arrow and a yellow dotted circle. Right: confocal maximum intensity projection showing input cells expressing GCaMP6f in the MEC. The white dotted line illustrates the impression made by the prism on the MEC. DG: dentate gyrus, Sub: subiculum. **B)** In vivo imaging example. Top left: imaging is performed with head-fixed mice under a two-photon benchtop microscope. Right: maximum intensity projection (across a 2-minute recording) of five cells in the MEC that provide monosynaptic input to the ipsilateral hippocampus 19 days after the rabies injection. Bottom left: ΔF/F traces of two cells with different activity onsets highlighted with corresponding colours in the image to the right.

### *In vivo* two-photon calcium imaging enables functional detection of distant input cells to birthdate-defined cell populations

While optogenetics is a powerful way to identify and characterise cells that are monosynaptically connected to birthdate-defined cell populations in the hippocampus, this approach is limited by the number of cells that can be recorded, and stimulated, at the same time. This is especially true if the labelled input cells are anatomically sparse. This becomes a pertinent issue when using rabies viruses, as it usually takes days or weeks to position tetrodes and long-term expression of rabies viruses will eventually lead to deterioration of cell health, which hence limit the time window for locating cells. We therefore tested the possibility of using calcium imaging to see if a rabies virus carrying GCaMP (RABV-GCaMP6f) injected into the hippocampus could be used to identify several input cells simultaneously in the MEC. We employed a similar approach to that described for the optogenetic experiments, except we implanted a prism between the MEC and cerebellum to obtain optical access to the superficial layers of MEC (**Figure 6A**). This enabled us to perform two-photon imaging in awake, head fixed mice and identify several cells projecting to a population of birthdate-matched cells in the hippocampus (**Figure 6B**).

Finally, to extend the time window for imaging experiments and ensure that the calcium signal itself is not altered by the overexpression of rabies proteins in the cell, we considered an alternative approach. By first introducing the calcium indicator with a separate AAV in the MEC of Cre and helper virus injected animals, dense population labelling can be achieved and imaging can be performed for as long as required. Then, a rabies virus carrying a red fluorescent tag can be injected into the hippocampus to identify which of the imaged cells project there, before the brain is preserved. This also enables activity profiles of targeted input cells to be compared to neighbouring cells (**Figure 7A**). For these reasons, this approach may be especially useful when attempting to identify inputs to a small number of starter cells, to help ensure their identification before their health deteriorates due to rabies virus overexpression. While following our protocol for targeting a single hippocampal starter cell, we injected an AAV carrying GCaMP6m (AAV-Syn-GCaMP6m) into the MEC and implanted a GRIN lens+prism doublet within the same surgery as the helper virus injection (**Figure 7A**). By keeping the craniotomy used for the helper virus injection free from cement and temporarily closing it with a silicone sealant, we could reopen it several weeks later to introduce the rabies virus carrying tdTomato (RABV-tdTomato). Once the rabies virus reached the MEC, we were able to identify a cell expressing both the calcium indicator and tdTomato *in vivo* with a two-photon miniscope (**Figure 7B**, Zong *et al*., 2017, 2021; Obenhaus *et al*., 2021), demonstrating the feasibility of using this approach to characterise the spatial firing properties of input cells in freely moving mice.

**Figure 7:**
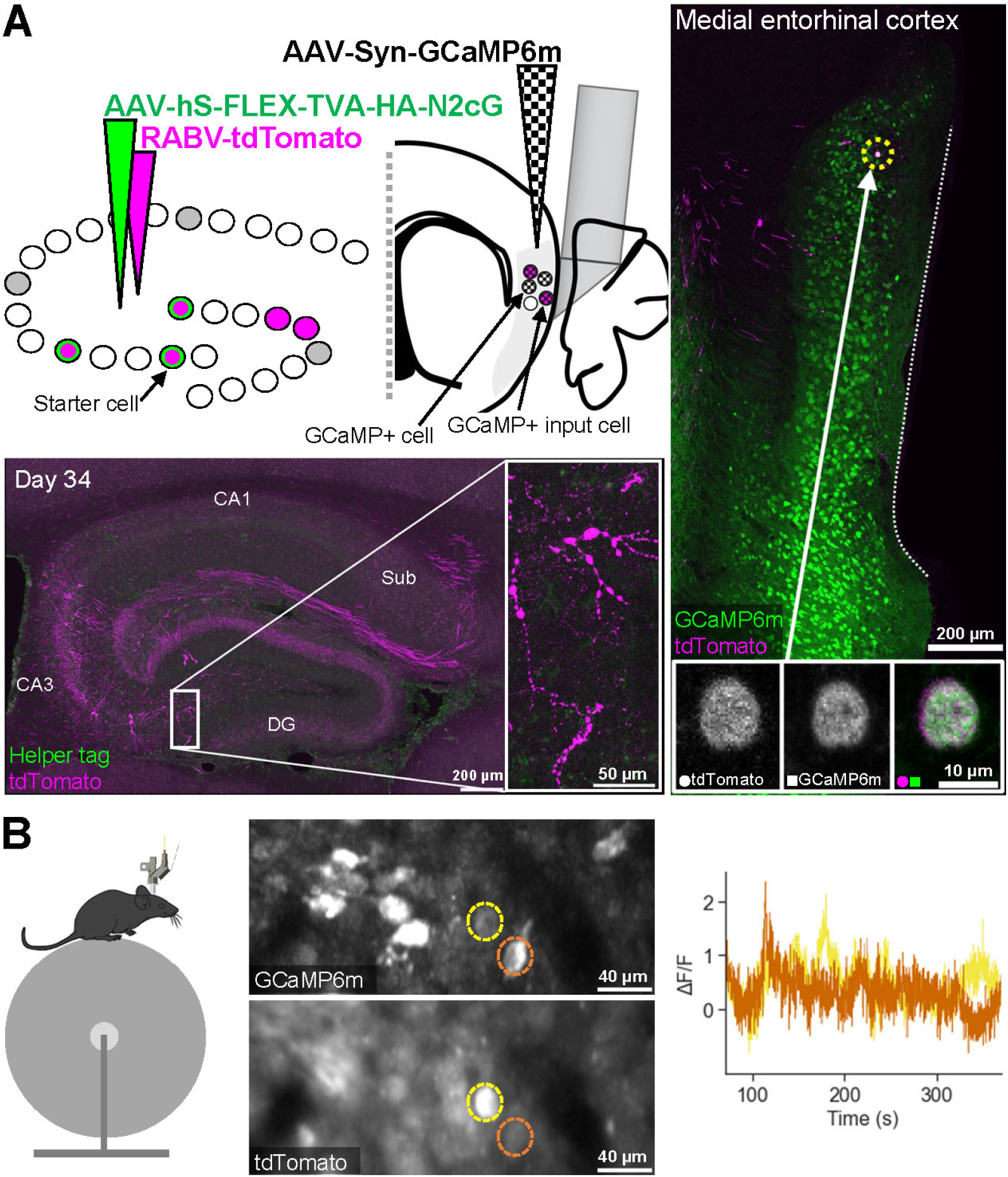
A dual labelling strategy facilitating long-term two-photon calcium imaging of distant input cells to birthdate-defined cell populations in vivo. **A)** Schematic of experimental design and resulting histology 34 days after the rabies injection. Top left: a population of birthdate-matched (E13) starter cells in the dorsal hippocampus is targeted to express TVA and G using the method described in Figure 1 (i.e. the number of starter cells is smaller than in Figure 6). In addition, an AAV virus carrying GCaMP6m is injected into the MEC (brain region marked in grey) for ubiquitous expression of the calcium indicator. A rabies virus carrying tdTomato injected into the hippocampus and a GRIN lens with a prism attached implanted by the MEC can then be used to identify cells in the MEC that provide monosynaptic input to the hippocampus. This approach enables cells providing input to the targeted starter cells and neighbouring cells to be compared, both before (avoiding the issue of rabies-mediated toxicity) and after rabies has been introduced. Bottom left: confocal maximum intensity projection of the dorsal hippocampus showing cells expressing tdTomato. The image inset shows a magnification of the region indicated by the white box, showing dendritic blebbing 34 days after the rabies injection. Right: confocal maximum intensity projection showing general GCaMP6m expression in the MEC and hippocampal input cells expressing tdTomato. Image insets are single z-plane confocal images showing colocalization of both markers in one example cell (i.e. an input cell also expressing GCaMP6m, highlighted with an arrow and yellow dotted circle). The white dotted line shows the impression made by the GRIN lens and prism on the MEC. DG: dentate gyrus, Sub: subiculum. **B)** In vivo imaging example. Left: imaging is performed in head-fixed mice with a two-photon miniscope. Middle: maximum intensity projection and mean projection (over time) of the GCaMP6m and tdTomato channels, respectively, acquired 29 days after the rabies injection. Both cells express GCaMP6m, but only one expresses tdTomato, meaning it provides monosynaptic input to the starter cell(s) targeted in the hippocampus. Right: ΔF/F GCaMP6m traces of the two cells highlighted with corresponding colours to the left.

## Discussion

We have developed a virus-based method that allows individual neurons to be targeted for monosynaptic rabies tracing in deep brain structures without the need for tissue aspiration. By applying this method to the hippocampus, we identify both local and distant inputs to single cells. We also demonstrate how this method can be used to determine the functional properties of monosynaptic inputs to birthdate-defined cell populations with *in vivo* electrophysiology combined with optogenetic tagging, or two-photon calcium imaging.

The present method achieves a success rate similar to that of existing approaches (few studies report overall success rate, but the 12% (successful transsynaptic spread in 14/119 electroporated cells) attained by Marshel *et al*. (2010) seems representative). However, it also offers several additional and unique advantages. First, it allows for rabies-mediated circuit mapping to be extended to single cells in sub-neocortical regions without the need for removal of overlaying tissue. Second, the *in utero* viral injection approach enables cells born at specific times to be targeted and hence the identification of inputs to cells with different developmental origins to be compared. Third, the use of viruses allows specific cell types, dividing cells or clones to be targeted by utilising cell-type specific gene regulatory elements (Nair *et al*., 2020) or different virus types (such as retroviruses), either during embryonic development (with the Cre-carrying virus) or in adult animals (TVA and G-carrying helper virus). Fourth, by simply varying the concentration and/or amount of virus used, any number of cells can be targeted for input mapping, from single neurons to populations of cells. Fifth, it can be used in wildtype animals, circumventing the need to generate and breed transgenic lines. Sixth, in theory this approach can be used in any species in which these viruses express and *in utero*/early postnatal injections are possible.

However, although modified rabies viruses are currently the only tools available for large scale monosynaptic input tracing *in vivo*, they also have disadvantages that need to be considered. Specifically, while the CVS-N2c strain used here is less toxic than the SAD-B19 strain (Reardon *et al*., 2016), it does eventually lead to death of the infected cells — a constraint it shares with all rabies virus-based approaches regardless of how cells are targeted. While the virus needs time to spread to distant input regions (e.g., to the MEC from the hippocampus, **Figure S2B**), waiting too long will result in cytotoxicity of the initially infected cells (**Figure 7A**) and hence an inability to determine the number of starter cells. From our experience, it appears that the rate at which this occurs differs based on the concentration and amount of virus the cells are exposed to (Lavin *et al*., 2020), their cell type, and the circuit that targeted cells are embedded in. However, with recent bioengineering efforts the limitation of rabies-mediated cytotoxicity can now be overcome, and thus long-term functional studies enabled, although additional viruses and/or transgenic animals are required to achieve this (Ciabatti *et al*., 2017, 2020; Chatterjee *et al*., 2018). Another limitation, also independent of the targeting approach, is the lack of knowledge regarding how rabies viruses spread to input cells. The number of input cells that project to a single cell remains subject to estimates but is likely to be substantially higher than what has been reported in single-cell rabies tracing studies (typically around 100 input cells and up to 846 in visual cortex (Wertz *et al*., 2015)). While time is a major determinant for how many input cells get labelled, the relatively low number typically seen raises the question of whether the virus exhibits a bias to cross synapses of certain types, strengths, or activity profiles. This remains an open question because a detailed ‘ground truth’ connectivity map, to which rabies tracing results can be compared, does not yet exist. The question of whether the transmission of rabies viruses is strictly synapse-specific is itself also under debate. It is therefore advisable to use rabies virus tracing for comparative and explorative studies rather than aiming for a complete quantification of all monosynaptic inputs to cells (see Rogers and Beier (2021) for a comprehensive review of these limitations).

The success rate of our method is chiefly determined by the ability to achieve satisfactory labelling of starter cells. In roughly half of the animals (**Figure 2A**), we could not determine how many starter cells were present due to ambiguous antibody staining of helper tags. The reason for this could be due to 1) death of starter cells or 2) staining issues (e.g., limited Cre availability resulting in few recombination events and/or inadequate amounts of helper proteins for our detection method). The latter explanation seems more likely for several reasons. First, we typically did not see cell damage until more than 30 days had passed after the rabies injection (**Figure 7A**). Second, we limited the amount of Cre and helper virus used to keep the number of starter cells low, which could result in reduced transgene expression from the helper virus. Third, animals that went through the same protocol, but were not injected with the rabies virus, also showed inconclusive staining of helper tags (n = 4 animals, data not shown). Fourth, antibody staining for helper proteins is clearer and easier to detect in animals injected with larger amounts of Cre and helper virus (**Figure 5, 6, S3**). Fifth, the spread of the rabies virus over time followed a similar trend for both the ‘inconclusive’ and ‘conclusive’ cases (**Figure 2B, S2**), indicating that the overall number of starter cells were similar in both groups. Sixth, based on the relatively low number of input cells observed, it seems unlikely that we are underestimating the number of starter cells. If anything, based on the low number of input cells we see, the number of starter cells should be even lower than what we report. Finally, antibody staining to detect starter cells is known to be challenging, because highly efficient interactions between the EnvA-pseudotyped rabies virus and its receptor TVA enable minimal expression of the latter for infection (Federspiel *et al*., 1994; Seidler *et al*., 2008). As a result, direct detection of the receptor or associated protein expression for starter cell identification can be difficult to achieve (Hafner *et al*., 2019; Lavin *et al*., 2020; Faulkner *et al*., 2021). Development of more sensitive and/or novel antibodies, for example against TVA or G specifically, will therefore likely improve the success rate of the present method and rabies tracing studies in general.

Rabies-mediated tracing from single cells is a powerful technique to map monosynaptic inputs, which is necessary to understand how neurons integrate information. With the present *in utero*-based virus injection method, rabies tracing can be achieved from individual cells located far beneath the brain’s surface. It thus extends detailed circuit mapping and the functional characterisation of local and distant inputs to deeper brain structures and to cells with defined developmental origins.

## Limitations of Study

While our method can target single cells in sub-neocortical regions for input mapping, it relies on rabies virus-mediated tracing (see Discussion for limitations) and *in utero* ventricular injections. The latter limits the cells that can be targeted to those that line the ventricles at the time of the injection. Even though that does include cortical cells types, existing methods such as single-cell electroporation (Marshel *et al*., 2010; Wertz *et al*., 2015; Rompani *et al*., 2017; Rossi, Harris and Carandini, 2020), patch clamping (Rancz *et al*., 2011; Vélez-Fort *et al*., 2014) or stamping (Schubert *et al*., 2017) approaches may be preferable. This is because reaching single cells in more readily accessible cortical areas involves minimal tissue damage and does not require embryonic injections. Our *in utero* protocol does, however, also enable our method to be used for developmental studies.

## Supporting information

Key resources table

Supplemental figures with legends

Supplemental tables with legends

## Acknowledgements

We thank S. Ball, A.M. Amundsgård, K. Haugen, K. Jenssen, E. Kråkvik, I. Ulsaker-Janke, and H. Waade for technical assistance, M.P. Witter for help identifying cell types and C. Kentros for discussions regarding the experimental protocol. We are also grateful to Tom M. Jessell and his lab at Columbia University for gifting us the essential reagents and cell lines to produce CVS-N2c rabies strain viruses when the project started in 2016. We are also thankful to Weijian Zong for providing us with a two-photon miniscope for recording GCaMP fluorescent activity in rabies virus injected mice. In addition, we want to thank Scidraw.io for making high quality drawings available, including contributions by Ann Kennedy of a mouse profile (https://doi.org/10.5281/zenodo.3925921).

This work was supported by a GRIDCODE ERC Advanced Investigator grant to E.I.M. (grant number 338865), the Research Council of Norway (RCN Centre of Excellence grant to M-B.M. and E.I.M., grant number 223262), an RCN infrastructure grant to E.I.M. and M-B.M. (NORBRAIN, grant number 295721), and the Kavli Foundation (M-B.M. and E.I.M.).

## Author contributions

Conceptualisation, R.I.J., F.D., M-B.M and E.I.M.; Methodology, R.I.J., R.R.N. and F.D.; Software, H.A.O.; Investigation, R.I.J., H.A.O., F.D., and T.S.; Resources, R.R.N.; Writing – Original Draft, R.I.J.; Writing – Review & Editing, R.I.J., R.R.N., H.A.O, F.D., T.S., M-B.M. and E.I.M.; Visualisation, R.I.J. and R.R.N.; Supervision, M-B.M. and E.I.M.; Funding Acquisition, M-B.M. and E.I.M.

## Inclusion and diversity statement

We worked to ensure sex balance in the selection of non-human subjects. One or more of the authors of this paper self-identifies as a member of the LGBTQ+ community.

## STAR Methods

### Resource availability

#### Lead contact

Further information and requests for resources and reagents should be directed to and will be fulfilled by the lead contact, Edvard I Moser.

#### Materials availability

The helper and rabies viruses used in this paper are available from the Viral Vector Core at the Kavli Institute for Systems Neuroscience, Norwegian University of Science and Technology (NTNU) in Trondheim, Norway.

#### Data and code availability

- Electrophysiology data used in this manuscript will be deposited at a repository and made publicly available as of the date of publication. The DOI will be listed in the key resources table.
- This paper does not report original code.
- Microscopy data and any additional information required to reanalyse the data reported in this paper is available from the lead contact upon request.

### Experimental model and subject details

Experiments were performed in accordance with the Norwegian Animal Welfare Act and the European Convention for the Protection of Vertebrate Animals used for Experimental and Other Scientific Purposes (permit numbers 6021, 7163 and 18013). C57BL/6JBomTac (Cat#B6JBOM; RRID: IMSR_TAC:b6jbom, Taconic) mice were housed in temperature- and humidity-controlled cages with sex-matched siblings when possible with food and water provided ad libitum. A reversed 12h light/12h darkness schedule was maintained throughout. Both sexes were used, which did not seem to influence the results (**Table S1, S2**), at embryonic and/or adult developmental stages.

### Method details

#### Helper virus (AAV-hS-FLEX-TVA-HA-N2cG)

A single Cre dependent rabies helper AAV that expresses the avian TVA receptor, 2xHA and rabies CVS-N2cG, under the human synapsin promoter, was designed and synthesized for the monosynaptic tracing experiments. The construct pAAV-hS-FLEX-splitTVA-2A-2xHA-2A-CVS-N2cG was created by replacing EGFP-B19G genes in pAAV-hSyn-FLEX-splitTVA-EGFP-B19G (Cat#52473, Addgene) by a 2xHA-CVS-N2cG sequence. The positive clones were confirmed by restriction digestion analyses and sequencing. Endotoxin free plasmid maxipreps (Cat#12663, Qiagen) were made for AAV preparations. The day before transfection, 7 × 10^6^ AAV-293 cells (Cat#CVCL_6871, Agilent, USA) were seeded in Dulbecco’s Modified Eagle Medium (DMEM, Cat#41965062, Thermo Fisher Scientific) containing 10% fetal bovine serum (FBS, Cat#16000-044, Thermo Fisher Scientific) and penicillin/streptomycin antibiotics (Cat#15140122, Thermo Fisher Scientific) into 150 mm cell culture plates. Calcium chloride mediated co-transfection was done with 22.5 µg pAAV containing the transgenes, 22.5 µg pHelper, 11.3 µg pRC (Cat#240071, Agilent, USA) and 11.3 µg pXR1 capsid plasmid (NGVB, IU, USA). After 7 hours, the medium was replaced with fresh 10% FBS containing DMEM. The transfected cells were scraped out after 72 hrs, centrifuged at 200 g and the cell pellet was subjected to lysis using 150 mM NaCl-20 mM Tris pH 8.0 buffer containing 10% sodium deoxy cholate. The lysate was then treated with Benzonase nuclease HC (Cat#71206-3, Millipore) for 45 minutes at 37 °C. The Benzonase treated lysate was centrifuged at 3000 g for 15 mins and the clear supernatant was then subjected to HiTrap® Heparin High Performance (Cat#17-0406-01, GE) affinity column chromatography using a peristaltic pump (McClure *et al*., 2011). The elute from the Heparin column was then concentrated using Amicon Ultra centrifugal filters (Cat# Z648043, Millipore). The titre of the viral stock was determined as approximately 10^11^ infectious particles/ml.

#### Rabies virus (RABV-GOI)

EnvA-pseudotyped G-deleted CVS-N2c rabies viruses expressing transgenes of interest (EnvA-ΔG-CVS-N2c RABV-GOI) were produced based on previous protocols (Wickersham, Sullivan and Seung, 2010; Osakada and Callaway, 2013). G-coated ΔG-CVS-N2c rabies viruses expressing transgenes of interest (ΔG-CVS-N2c RV-GOI), supernatants and packaging cell lines were kind gifts from the Jessell lab (Reardon *et al*., 2016). The uninfected packaging cells were maintained in 10% FBS and Gentamicin (Cat#15750060, ThermoFisher Scientific) containing EMEM (Eagle’s Minimal Essential Medium, Cat#30-2003, ATCC) in a humidified atmosphere of 5% CO_2_ at 37°C. ΔG-CVS-N2c RV-GOI viruses were amplified by passaging for several days on N2cG-complementing Neuro2A-N2c(G) cells. During this expansion process, the infected cells were grown in humidified atmosphere of 3% CO_2_ at 34°C. The supernatants were harvested, filter sterilized and frozen every three days for 4-6 weeks. The functional titre of various batches of ΔG-CVS-N2c RV-GOI was determined by infecting Neuro2A cells (Cat#CCL-131, ATCC) and quantifying the GOI positive Neuro2A cells. For pseudotyping with EnvA, Neuro2A-EnvA cells were infected with unpseudotyped ΔG-CVS-N2c RV-GOI particles at a multiplicity of infection less than 0.5. Twenty-four hours post-infection, infected cells were washed with DPBS (Dulbecco’s phosphate buffered solution, Cat#14190169, ThermoFisher Scientific), trypsinized with 0.25% trypsin-EDTA (Cat#T4049-500ML, Merck) and reseeded on multiple 150 mm dishes in a humidified atmosphere of 3% CO_2_ at 34°C. For producing high titre EnvA-pseudotyped ΔG-CVS-N2c RABV-GOIs, incubation media was harvested 48 hrs later, filtered with 0.45 µm filter and viral particles were concentrated by ultracentrifugation at approximately 50,000 g for 2 hrs at 4°C. The pellets were resuspended in DPBS and concentrated using Amicon Ultra centrifugal filters (Cat#Z648043, Millipore). EnvA-pseudotyped ΔG-CVS-N2c rabies was titrated by serial dilutions with HEK293-TVA800 cells. The titre of the viral stock (EnvA-ΔG-CVS-N2c RABV-GOI) was determined as approximately 10^7^ infectious particles/ml.

#### *In utero* injections

Two females and one male were put in the same cage for 24 hours. Embryonic day 1 (E1) was defined as the day on which the male was removed. Females that were suspected to be pregnant were anaesthetised with isoflurane (IsoFlo, Zoetis, 5 % isoflurane vaporised in medical air delivered at 0.8-1 l/min for induction, then 1.5-3 %) and the fur covering the abdomen was removed with hair removal cream for sensitive skin (Veet, Canada). If pregnancy was confirmed with ultrasound (Vevo 1100 System with MS-550S probe, Fujifilm Visualsonics) the mouse was prepared for surgery by injecting an analgesic subcutaneously (Metacam, Boehringer Ingelheim, 5 mg/kg) and placing her on a heated surgery stage (37°C). Midline incisions were made in the skin and peritoneum, and warm (37-39 °C) saline (NaCl 9 mg/ml, B. Braun Medical) was used throughout to keep the abdominal cavity warm and wet. Embryos were carefully transferred either one by one or in pairs to a decontaminated chamber placed on top of the female’s abdomen and covered with warm ultrasound gel (37-39 °C, Aquasonic 100, Parker). One lateral ventricle in all embryos in both uterine horns were injected with AAV-CKII-Cre (AAV1.CamKII0.4.Cre.SV40, Cat#AV-1-PV2396, titre: 2.71e13 GC/ml, UPenn Vector Core, University of Pennsylvania, USA), diluted when applicable (see **Table S1, S2**) in sterile DPBS (1X Dulbecco’s Phosphate Buffered Saline, Gibco, ThermoFisher), with a pressure-pump (Visualsonics) under ultrasound guidance. Custom made glass micropipettes (outer tip opening: 200 μm, inner tip opening: 50 µm, Origio) were used to inject the virus in six pulses of 50.6 nl each. After the pipette had been retracted, the embryos were wiped clean with sterile gauze and gently returned to the abdominal cavity. Once injections were complete, the peritoneum and skin were sutured (Supramid DS 13, Resorba Medical, Germany) separately. The entire procedure, from first incision to last stitch, was limited to maximum 90 min to minimise stress to the animals. Recovery took place in a heated chamber (33°C) until mobility and alertness was restored.

#### Adult injections

Once *in utero* injected embryos had reached adulthood (minimum 90 days after birth), they were prepared for the second virus injection. After being anaesthetised with isoflurane (IsoFlo, Zoetis, 5 % isoflurane vaporised in medical air delivered at 0.8-1 l/min) two analgesics were provided subcutaneously (Metacam, Boehringer Ingelheim, 5 mg/kg or Rimadyl, Pfizer, 5 mg/kg, and Temgesic, Indivior, 0.05-0.1 mg/kg) and one local anaesthetic underneath the skin covering the skull (Marcain, Aspen, 1-3 mg/kg). The fur on the top of the head was then shaved and the mouse placed in a stereotaxic frame, the isoflurane level adjusted to 1-2 % and eye ointment applied to prevent dry eyes (Simplex, Actavis). Next the skull was exposed and a small hole drilled (with Cat#1RF HP 330 104 001 001 005 drill bit from Hagen and Meisinger, Germany) at 1.55 mm lateral from the midline and 1.58 mm posterior from bregma in the right hemisphere. The same coordinates were used to inject the helper virus (AAV-hS-FLEX-TVA-HA-N2cG, made in house) into the dorsal hippocampus at 1.55 mm depth from the dura with a pressure-pump (Nanoject, Visualsonics) and pipettes (Cat#504949, WPI) pulled and cut to produce an extended tip with 20-50 µm outer diameter. The required volume (see **Table S1, S2**) was injected at a rate of 23 nl/sec and the pipette left for 10 min before retracting. The exposed brain was kept wet with saline (NaCl 9 mg/ml, B. Braun Medical) throughout. The craniotomy was then covered with UV curable cement (Venus Diamond Flow, Kulzer) and the skin sutured (Supramid DS 13, Resorba Medical, Germany). The third virus injection was made at least two weeks later using the same procedure (after removing the cement-cover with forceps and reopening the craniotomy with a drill if necessary) except 180-560 nl of the modified rabies virus (EnvA-pseudotyped ΔG-CVS-N2c RABV-GOI, see “Rabies virus (RABV-GOI)”) was injected across 1.7-1.40 mm below the dura over at least 10 min. Recovery took place in a heated chamber (33°C) until mobility and alertness was restored. A second subcutaneous injection of the analgesic Metacam (Boehringer Ingelheim, 5 mg/kg) or Rimadyl (Pfizer, 5 mg/kg) was provided under isoflurane anaesthesia once per day for 1-2 days after surgery.

Animals not following the ‘single cell experiment’ protocol underwent the same procedure but modified to suit the particular experiment as indicated in the text, figure legends and/or **Table S2** (e.g. only one injection and/or different viruses, including AAV1-CAG-FLEX-tdTomato (titre: 5.88e12 GC/ml, Cat#AV-1-ALL864, UPenn Vector Core, University of Pennsylvania, USA) in **Figure S1B**).

#### Optrode construction

Four 17 mm polyimide-coated platinum-iridium (90–10%) wires (California Fine Wire, CA) were twisted to create one tetrode. Four tetrodes were wired to a 16 channel microdrive (Axona Ltd., Herts, UK) via a connected 22G metal cannula encased in an 18G outer cannula (with one end cut and polished smooth to accommodate the brain surface curvature during implantation) and cut to length with sharp scissors. A 100 µm in diameter optic fibre (Cat#MFC_100/125–0.37_17 mm_ZF1.25-C45, Doric Lenses) was glued to the anterior side of the tetrodes with the tip approximately 200 µm above the tetrode tips. Electrode impedance was reduced to between 183 and 220 kOhm by plating the electrode tips with platinum at 1 kHz and -0.17 µA using a NanoZ device (Neuralynx, Bozeman, USA).

#### Optrode implantation

One E12 *in utero* AAV-CKII-Cre (no dilution) injected mouse and one that had the Cre virus introduced with the AAV-hS-FLEX-TVA-HA-N2cG virus (mixed 1:1, total volume 303.6 nl) at the adult stage were used. Virus injections were performed as described under “Adult injections” with the rabies virus carrying ChR2 and YFP (RABV-ChR2-YFP). An optrode was also implanted in the MEC in the ipsilateral (right) hemisphere in the same surgery as the rabies injection as follows: after exposing and scratching the skull with a needle to enhance skull-dental cement adherence, one ground screw with a short wire soldered on (which was then soldered to the optrode at the end/after the surgery) was attached to the occipital bone and one extra screw to the parietal bone, both in the left hemisphere. Tissue adhesive (Histoacryl, B. Braun Surgical) was applied to secure the screws and to attach the skin edges to the lateral edges of the skull. A circular craniotomy was made over and anterior to the transverse sinus in the right hemisphere. An area of the dura slightly bigger than the optrode to be implanted was removed anterior to the sinus. The optrode was implanted 3.5 mm lateral from the midline, 0.4 mm anterior to the sinus and at a 6 degree angle (pointing posteriorly) down to 0.9-1 mm depth below the dura. The outer cannula was gently lowered to touch the dura surrounding the implant. The implant was secured to the skull with UV curable dental cement (Venus Diamond Flow, Kulzer), and dental cement (Paladur, Kulzer) mixed with carbon (CAS number 7440-44-0, Sigma-Aldrich) to reduce transparency and light reflections. 0.5 ml saline (NaCl 9 mg/ml, B. Braun Medical) warmed to body temperature was administered 1-2 times during the procedure to maintain hydration. Recovery took place in a heated chamber (33°C) until mobility and alertness was restored. A second subcutaneous injection of the analgesic Temgesic (Indivior, 0.05-0.1 mg/kg) was provided approximately eight hours after the first and Metacam (Boehringer Ingelheim, 5 mg/kg) or Rimadyl (Pfizer, 5 mg/kg) was readministered once per day for at least two days after the surgery, all under isoflurane anaesthesia.

#### *In vivo* electrophysiology

Animals implanted with an optrode were recorded up to two times per day. Before each recording session the microdrive was connected to a digital acquisition system (Axona) to record spikes, and two LEDs attached to the drive to enable tracking of position and head direction with a tracking system (Axona). Signals from the tetrodes were amplified 5 000-10 000 times and a 0.3-7 kHz bandpass filter was applied. Individual spikes were stored at 48 kHz (8 bits/sample and 50 samples per waveform) with a 32-bit time stamp (96 kHz clock rate). LED position data were recorded at 50 Hz. Each session consisted of two parts. First, the mouse explored an open field environment (1 × 1 ×0.5 m black box with a white, rectangular cue card fixed to one of its walls) for approximately 30 min. Cookie crumbs were thrown into the box throughout until the mice were satisfied and stopped eating. Second, the mouse was placed in a plexiglass holding box (20 × 25 × 15 cm, lined with a cotton towel) outside and above the open field arena. In this second part, a 473 nm laser (Cobolt 06-MLD, output power max: 100 mW, Hübner Photonics) was connected to the implanted optic fibre via a patch cable (Doric Lenses, Canada) after the power had been adjusted to deliver 7.5-15.14 mW at the end of the patch cable. TTL pulses sent from an Arduino Uno Rev3 (Arduino, Italy) controlled the on/off state of the laser to deliver 1 ms light pulses at 10 Hz continuously. The same TTL was fed in parallel into a spare digital input channel of the Axona acquisition system for precise synchronisation of laser stimuli and electrophysiological data. At the end of each recording session the tetrodes were turned down by 25-50 µm (except when trying to record from the same cell across multiple sessions, **Figure S3**). To encourage exploration in the open field environment, animals were put on a food restriction schedule outside of the recording sessions. Their weight was monitored daily and on average was maintained at 96% of their baseline weight.

#### Prism implantation with RABV-GCaMP6f

To visualise distant inputs located in the MEC *in vivo*, the protocol illustrated in **Figure 1** was modified as follows: during the *in utero* injection (as described under “*In utero* injections”), the AAV-CKII-Cre virus was not diluted and when the AAV-hS-FLEX-TVA-HA-N2cG virus was injected (as described under “Adult injections”) a larger volume was used (**Table S2**). This resulted in a large population of starter cells in the hippocampus. In the same surgery as the rabies virus injection (as described under “Adult injections” except here RABV-GCaMP6f was used, see “Rabies virus (RABV-GOI)”), the skull was exposed, the rabies virus injected, and tissue adhesive (Histoacryl, B. Braun Surgical) was applied to attach the skin edges to the lateral edges of the skull. Next, the skull was scratched with a needle to enhance skull-dental cement adherence and a large, circular craniotomy over and behind the transverse sinus in the right hemisphere was made. The dura connecting the sinus to the cerebellum was carefully cut along the posterior side of the sinus before a 2 mm square microprism with reflective enhanced aluminium coating on the hypotenuse (Tower Optical, USA) glued (Norland Optical Adhesive 71, Norland Products) to a 4 mm in diameter circular cover slip (64-0724, Warner Instruments) was inserted immediately posterior to the sinus by hand. No brain tissue was removed. A thin, metal cannula attached to the stereotaxic frame was used to hold the prism in place after insertion while the coverslip was secured to the skull with UV curable cement (Venus Diamond Flow, Kulzer). Dental cement (Paladur, Kulzer) mixed with carbon (CAS number 7440-44-0, Sigma-Aldrich; to reduce transparency and light reflections) was used to attach a custom-made head bar and to cover the rest of the skull. 0.9 ml saline (NaCl 9 mg/ml, B. Braun Medical) warmed to body temperature was administered subcutaneously at the end of the procedure for hydration. Recovery took place in a heated chamber (33°C) until mobility and alertness was restored. A second subcutaneous injection of the analgesic Temgesic (Indivior, 0.05-0.1 mg/kg) was provided approximately eight hours after the first and Rimadyl (Pfizer, 5 mg/kg) was readministered once per day for at least two days after surgery, all under isoflurane anaesthesia.

#### Lens+prism implantation with RABV-tdTomato

One mouse went through the same ‘single cell experiment’ protocol illustrated in **Figure 1** but adapted to enable imaging of distant input cells in the MEC as follows: in the surgery when the AAV-hS-FLEX-TVA-HA-N2cG virus was injected (as described under “Adult injections”), the craniotomy over the hippocampus was not covered but encircled with UV curable cement (Venus Diamond Flow, Kulzer) to create a small well that was filled with a silicone sealant (Kwik-Cast, World Precision Instruments). In addition, tissue adhesive (Histoacryl, B. Braun Surgical) was applied to attach the skin edges to the lateral edges of the skull. Next, a large, oval craniotomy centred over the transverse sinus in the right hemisphere was made. Here, two injections of an AAV-Syn-GCaMP6m virus (AAV1.Syn.GCaMP6m.WPRE.SV40, titre: 3.43e13 GC/ml, Cat#AV-1-PV2823, UPenn Vector Core, University of Pennsylvania, USA) was performed at 3.5 mm and 3.9 mm lateral from the midline, 0.2 mm anterior to the sinus and at a 7 degree angle (pointing posteriorly) with a microliter syringe (#75, Hamilton Company). The syringe was slowly lowered until it touched the dura covering the MEC (typically after approximately 2 mm). The syringe was then raised by 100 µm before injecting 500 nl at 100 nl/min across 400 µm for each injection. After each injection, the syringe was left in place for 10 min before retracting. The craniotomy was then temporarily covered with a silicone sealant (Kwik-Cast, World Precision Instruments) while a bonding agent (OptiBond All-In-One, Kerr) was applied to the skull. A custom-made head bar was next attached in front of the hippocampus craniotomy with dental cement (Paladur, Kulzer). Once the cement had dried, the ear bars stabilising the skull were removed to ease the pressure on the skull while the headbar ensured that the skull remained fixed in the same position. Next the silicone sealant covering the MEC craniotomy was removed and the dura connecting the sinus to the cerebellum was carefully cut along the length of the sinus with a needle. Then, a 1 mm in diameter GRIN (gradient-index) lens attached to a 1 mm square prism (total length approximately 4.67 mm, optimised for 920 nm, Grintech, Germany) was *slowly* inserted between the cerebral cortex and cerebellum at an 8 degree angle (pointing anteriorly) with a custom made holder attached to a stereotactic micromanipulator (Cat#1760, Kopf, USA). No brain tissue was removed. The GRIN lens+prism was positioned as laterally as possible while allowing for most of the prism itself to be immersed and simultaneously moved anteriorly to ensure contact with the surface of the MEC and thereby limit movement during *in vivo* imaging. Once in place, the craniotomy was covered with another silicone sealant (Kwik-Sil, World Precision Instruments) and a strip of UV curable cement (Venus Diamond Flow, Kulzer) applied to help maintain the sealant and implant in place while removing the lens-holder. The same cement was then used to encase the entire craniotomy and secure the implant before applying dental cement (Paladur, Kulzer) mixed with carbon (CAS number 7440-44-0, Sigma-Aldrich; to reduce transparency and light reflections) on top and over the rest of the skull (avoiding the hippocampus craniotomy). For protection a small piece of micropore tape (3M) was glued over the hippocampus craniotomy and the GRIN lens+prism doublet was covered with Kwik-Cast. 0.3 ml saline (NaCl 9 mg/ml, B. Braun Medical) was administered subcutaneously at the end of the procedure for hydration. Recovery took place in a heated chamber (33°C) until mobility and alertness was restored. A second subcutaneous injection of the analgesic Temgesic (Indivior, 0.05-0.1 mg/kg) was provided approximately eight hours after the first and Metacam (Boehringer Ingelheim, 5 mg/kg) was readministered once per day for at least two days after surgery, all under isoflurane anaesthesia. Three weeks later the mouse was again anaesthetised with isoflurane (IsoFlo, Zoetis, 5 % isoflurane vaporised in medical air delivered at 0.8-1 l/min) and one analgesic was provided subcutaneously (Metacam, Boehringer Ingelheim, 5 mg/kg) before reducing the isoflurane level to 1-2 % and applying eye ointment (Simplex, Actavis). The skull was realigned to the same position as the previous surgery using the head bar and the tape and silicone sealant covering the craniotomy over the hippocampus was removed. After reopening the craniotomy RABV-tdTomato was injected in the centre of the craniotomy but otherwise in the same way as described under “Adult injections” and lastly sealed with UV curable cement (Venus Diamond Flow, Kulzer).

#### *In vivo* two-photon calcium imaging

Mice were imaged with a custom two-photon benchtop microscope (Femtonics, Hungary) or two-photon miniscope (Zong *et al*., 2017, 2021; Obenhaus *et al*., 2021). In both cases, the mice were head-fixed via their head bar and allowed to run freely on a black styrofoam wheel (approximately 85 mm radius and 70 mm width) with a metal shaft fixed through the centre. Each end of the metal shaft was attached to a low friction ball bearing (HK 0608, Kulelager AS, Norway) that was held in place on the optical table with a custom mount (https://github.com/kavli-ntnu/wheel_tracker). On the benchtop microscope, a Ti:Sapphire laser (MaiTai Deepsee eHP DS, Spectra-Physics) tuned to 920 nm was transmitted through a 16x/0.8NA water-immersion objective (Cat#MRP07220, Nikon), after which its power was adjusted to measure <180 mW (using S121C connected to PM100D, Thorlabs, USA). Emitted fluorescence was routed to a GaAsP detector via a 600 nm dichroic mirror and 490-555 nm band pass filter supplied with the microscope. The MESc software (Femtonics, Hungary) accompanying the microscope was used for microscope control and image acquisition. 620 x 620 µm images were acquired at 512 x 512 pixel resolution with a frame rate of approximately 30 Hz.

For miniscope imaging, a custom made setup described in detail by Obenhaus et al. (2021) was used in conjunction with a Ti:Sapphire laser (MaiTai Deepsee eHP DS, Spectra-Physics) tuned to 920 nm to acquire dual-channel images at 512 x 512 pixel resolution with a frame rate of approximately 7.52 Hz.

#### Histology

Following induction of very deep anaesthesia with isoflurane and a 0.3 ml intraperitoneal injection of Pentobarbital (Apotekforeningen, 100mg/ml), mice were transcardially perfused with saline (0.9% NaCl in purified water) and then day-fresh 4 % PFA (paraformaldehyde, CAS Number: 30525-89-4, Alfa Aesar) in phosphate buffered saline (PBS, P3813, Sigma-Aldrich, Germany). The extracted brains were left in 4 % PFA at 4 °C for up to 24 hours before being transferred to a 30 % sucrose solution in PBS for at least 2 days. 40 µm sagittal sections were made on a cryostat (CryoStar NX70, Thermo Scientific, USA), taking care to maintain their sequential order in a 24-well plate filled with anti-freeze solution (40 % PBS, 30 % glycerol, 30 % ethylene glycol). The sections were stored in a –20 °C freezer. Before staining, sections were washed three times in PBS.

Sections from animals that went through the ‘single-cell experiment’ method illustrated in **Figure 1** were either stained using a standard protocol or a TSA Plus (Akoya Biosciences, USA) protocol as indicated in **Table S1** to label cells expressing the helper virus. The TSA Plus protocol was adopted in an attempt to improve labelling of starter cells, but both protocols yielded comparable results. Sections from other animals were stained with the standard protocol. During all washes and incubations (except the one at 37 °C) sections were left on a shaker. All washes were approximately 10 min long.

Standard protocol: three washes in a 0.3 % Triton X-100 (Merck) in PBS solution (‘PBS-T’) before incubation in a 3 % BSA solution (bovine serum albumin, Cat#a2153, CAS Number: 9048-46-8, Sigma; in PBS-T) with primary antibodies against HA (rabbit, Cat#3724, Cell Signaling Technology, USA; dilution 1:500) or 2A (rabbit, Cat#ABS31, Merck, Germany; dilution 1:500), and RFP (rat, Cat#5f8, ChromoTek, Germany; dilution 1:500) or GFP (chicken, Cat#ab13970, Abcam, UK; dilution 1:1000) as appropriate for two days at approximately 10 °C. After three washes in PBS-T, sections were again incubated in the 3 % BSA solution but with secondary antibodies against rabbit (with AF647 conjugate, Cat#A31573, Invitrogen; dilution matching the primary antibody), and rat (with AF555 conjugate, Cat#ab150166, Abcam, UK, or AF594 conjugate, Cat#A-21209 Invitrogen; dilution 1:500) or chicken (with AF488 conjugate, Cat#A11039, Invitrogen; dilution 1:1000) as appropriate for 3 hours at room temperature (RT).

They were then washed three times in PBS before mounting onto Polysine adhesion slides (Cat#10219280, Brand: J2800AMNZ, Thermo Scientific, US) in PBS and coverslipped with Prolong antifade with DAPI (Cat#P36935, Invitrogen). TSA Plus protocol: sections were first incubated with 0.01 % H_2_O_2_ in PBS at RT for 30 min before being washed three times in PBS. They were then incubated in a 3 % BSA solution containing primary antibodies as in the standard protocol (except a 1:1000 dilution was used for the antibody against HA) for two days at approximately 10 °C. After two washes in TNT (3.03 g Tris, 4.48 g NaCl, 2.5 ml Tween in 500 ml purified water, pH adjusted to 8 with HCl), they were incubated in TNB (TNT with blocking reagent (Cat#FP1012, Akoya Biosciences)) for 1 hr at RT before being transferred to TNB containing antibodies against rabbit (with HRP conjugate, goat, Cat#G-21234, Invitrogen, US; dilution 1:400) and rat (with AF555 conjugate, Cat#ab150166, Abcam, UK; dilution 1:1000) for 2 hours at 37 °C. Two washes in TNT was followed by an incubation in an amplification solution (Cat#FP1498, Akoya Biosciences) containing an antibody against HRP (with Cy5 conjugate, Cat#NEL745001KT, Akoya Biosciences; dilution 1:200) at RT for 30 min. The sections were then washed three times in TNT before mounting as in the standard protocol.

#### Microscopy of stained sections

All sections containing rabies labelled cells in the dorsal hippocampus and entorhinal cortex were first imaged with an epifluorescence microscope (AxioImager Z1, Zeiss, Germany) using a 10x/0.45NA objective (Cat#1063-139, Zeiss Plan-Apochromat, Zeiss, Germany) and manually checked and confirmed with higher magnifications. Individual rabies labelled cells were then scanned with a confocal microscope (LSM 880, Zeiss, Germany) using a 40x/1.4NA oil immersion objective (Cat#420762-9900, Zeiss Plan-Achromat, Zeiss, Germany) at multiple optical planes 0.5-1 µm apart to check for colocalization between the proteins introduced by the helper virus and that introduced by the rabies virus. Some sections were also scanned with a 20x/0.8NA air objective (Zeiss Plan-Achromat, Zeiss, Germany) to create overview images.

### Quantification and statistical analysis

Details of statistical tests can be found in the figure legends. Animals that went through the ‘single cell experiment’ protocol described in **Figure 1** were excluded if one or both adult virus injections missed the hippocampus (n = 2 animals) or a section containing the dorsal hippocampus was lost during histological processing (n = 1 animal). Analysis of electrophysiology and *in vivo* imaging data were integrated in a custom Datajoint framework (Yatsenko *et al*., 2015).

#### Conclusive/inconclusive classification

All rabies positive cells in the dorsal hippocampus were examined manually in Fiji (Schindelin *et al*., 2012) to check for colocalization with proteins introduced by the helper virus in confocal stacks acquired at 40x. Cells expressing both helper and rabies proteins were determined to be ‘conclusive’ starter cells if their labelling was unambiguous, i.e. the label was in the cell body, clearly distinguishable from background, and visible in more than one optical section. If no conclusive cells were found in an animal, it was considered ‘inconclusive’. Antibodies against 2A and HA (‘helper tags’) produced similar staining results and were treated equally during analysis.

#### Spike sorting

Spikes recorded from both the first and second part of an electrophysiological recording session were merged and sorted together offline with KlustaKwik (Kadir, Goodman and Harris, 2014; Rossant *et al*., 2016), first automatically and then refined manually (e.g. merging of clusters) in a graphical user interface (KlustaViewa). The respective recordings (open field and laser stimulation trials) were then split apart again for subsequent analysis.

#### Analysis of electrophysiology data

All data were analysed with custom Python code (Python Software Foundation) as reported in Rowland *et al*. (2018). Spike latencies in laser stimulation sessions was calculated as the average time between a laser pulse (a ‘trial’) and the first spike to occur within 10 ms (trials in which a spike did not occur within 10 ms were not included). Fidelity was calculated as the percentage of laser stimulation trials in which a spike occurred within 10 ms of the laser pulse. The head direction score and mean vector length for cells in **Figure S4** were calculated as described previously (Rowland *et al*., 2018).

#### Analysis of calcium imaging data

All *in vivo* acquired images were motion corrected and ROIs extracted using the Suite2p (Pachitariu *et al*., 2017) python library (https://github.com/MouseLand/suite2p). The Suite2p GUI was used to manually refine the ROI selection (e.g. discard obvious artefactual ROIs). ΔF/F was calculated by first multiplying the neuropil fluorescence of every cell with a neuropil coefficient (0.7) and subtracting the resultant trace from the raw fluorescence trace. The baseline for every cell was then subtracted and the result divided by baseline (Obenhaus *et al*., 2021; Zong *et al*., 2021). The baseline was calculated as the median of the first 20% of frames sorted ascendingly after smoothing the neuropil corrected trace with a hanning window.

