## Supplementary material for "All-viral tracing of monosynaptic inputs to single birthdate-defined neurons in the intact brain": Key resources table

| REAGENT or RESOURCE | SOURCE | IDENTIFIER |
| --- | --- | --- |
| **Antibodies** | | |
| Anti-HA, rabbit | Cell Signaling Technology | Cat#3724 |
| Anti-2A, rabbit | Merck | Cat#ABS31 |
| Anti-rabbit, AF647 | Invitrogen | Cat#A31573 |
| Anti-rabbit, HRP | Invitrogen | Cat#G-21234 |
| Anti-HRP, Cy5 | Akoya Biosciences | Cat#NEL745001KT |
| Anti-RFP, rat | ChromoTek | Cat#5f8 |
| Anti-rat, AF555 | Abcam | Cat#ab150166 |
| Anti-rat, AF594 | Invitrogen | Cat#A-21209 |
| Anti-GFP, chicken | Abcam | Cat#ab13970 |
| Anti-chicken, AF488 | Invitrogen | Cat#A11039 |
| **Bacterial and virus strains** | | |
| AAV1.CamKII0.4.Cre.SV40 (AAV-CKII-Cre) | UPenn Vector Core | Cat#AV-1-PV2396 |
| AAV-hS-FLEX-TVA-HA-N2cG | This paper / Viral Vector Core at Kavli Institute for Systems Neuroscience | N/A |
| EnvA-ΔG-CVS-N2c RV-tdTomato (RABV-tdTomato) | As above | N/A |
| EnvA-ΔG-CVS-N2c RV-ChR2-YFP (RABV-ChR2-YFP) | As above | N/A |
| EnvA-ΔG-CVS-N2c RV-GCaMP6f (RABV-GCaMP6f) | As above | N/A |
| AAV1.CAG.Flex.tdTomato.WPRE.bGH | UPenn Vector Core | Cat#AV-1-ALL864 |
| AAV1.Syn.GCaMP6m.WPRE.SV40 | UPenn Vector Core | Cat#AV-1-PV2823 |
| **Deposited data** | | |
| *In vivo* electrophysiology data | This paper | Will be added upon publication |
| **Experimental models: cell lines** | | |
| Neuro2A-CVS-N2c(G) | Jessell lab (Reardon *et al.*, 2016) | N/A |
| Neuro2A-EnvA | Jessell lab (Reardon *et al.*, 2016) | N/A |
| Neuro2A | American Type Culture Collection (ATCC) | CCL-131 |
| HEK293-TVA800 | Callaway lab (Osakada and Callaway, 2013) | N/A |
| **Experimental models: Organisms/strains** | | |
| Mice: C57BL/6JBomTac | Taconic | Cat#B6JBOM; RRID: IMSR_TAC:b6jbom |
| **Recombinant DNA** | | |
| pAAV-hSyn-FLEX-splitTVA-2A-2xHA-2A-CVS-N2cG | This paper / Viral Vector Core at Kavli Institute for Systems Neuroscience | N/A |
| **Software and algorithms** | | |
| KlustaKwik | Kadir, Goodman and Harris, 2014; Rossant *et al.*, 2016 | https://github.com/klusta-team/klustakwik |
| Python | Python Software Foundation | https://www.python.org/ |

| | | |
|---|---|---|
| Datajoint | Yatsenko *et al.*, 2015 | https://github.com/datajoint/datajoint-python |
| Suite2p python | Pachitariu *et al.*, 2017 | https://github.com/MouseLand/suite2p |
| Fiji (ImageJ) | Schindelin *et al.*, 2012 | http://fiji.sc; RRID: SCR_002285 |
