## Supplemental figures with legends for "All-viral tracing of monosynaptic inputs to single birthdate-defined neurons in the intact brain"

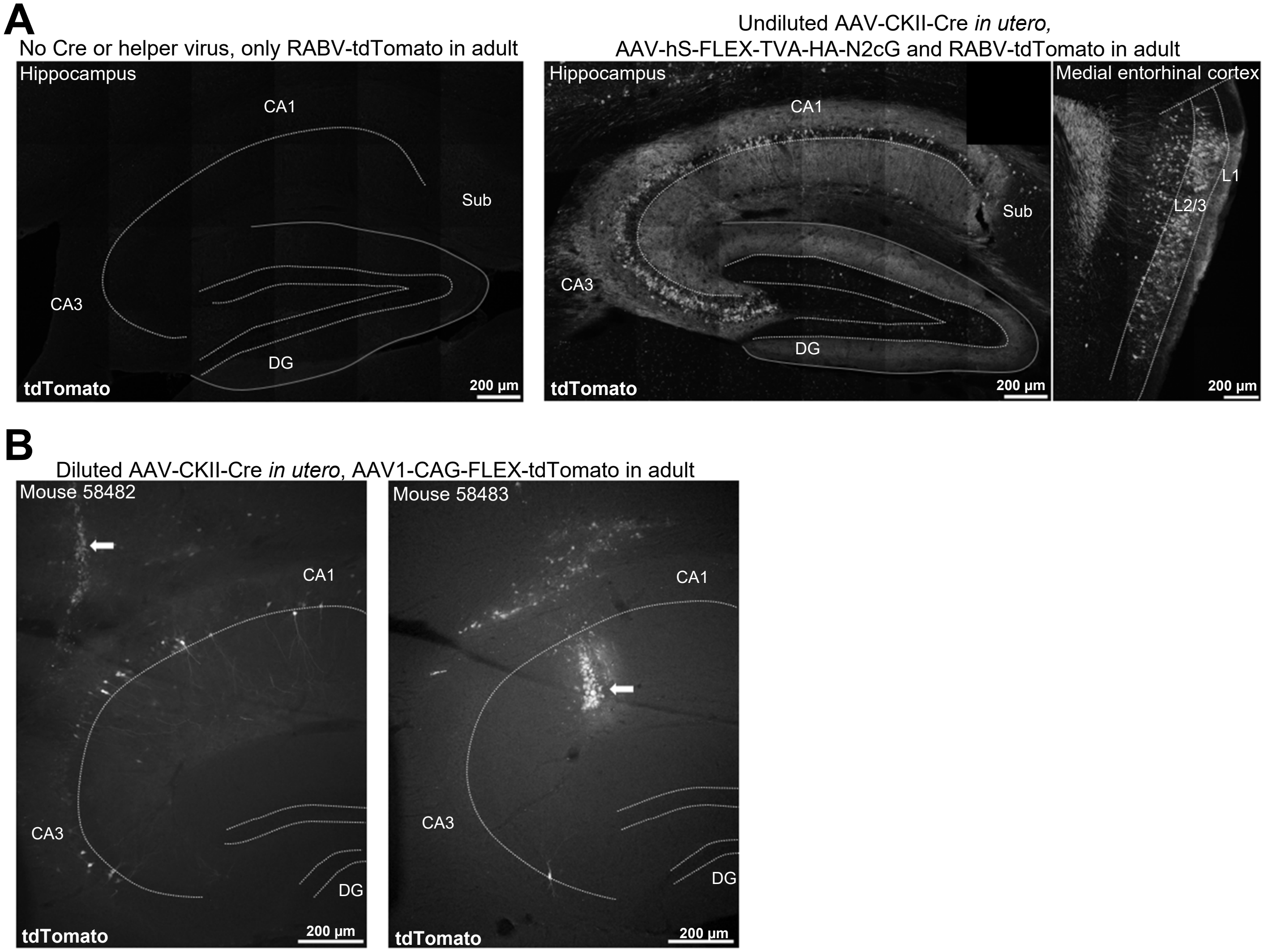


**Figure S1: No rabies expression without TVA and G, and confirmation of sparse Cre expression, related to figure 1**

**A)** Left: no tdTomato expression in a mouse in which only RABV-tdTomato was injected in the hippocampus. Right: extensive local and distant tdTomato expression in a mouse injected with undiluted AAV-CKII-Cre *in utero* at E12 followed by a large AAV-hS-FLEX-TVA-HA-N2cG injection and separate RABV-tdTomato injection in the adult hippocampus. Images are maximum intensity projections of confocal images. The amount of RABV-tdTomato injected was the same for both mice and they were both perfused 14 days after the rabies injection. DG: dentate gyrus, Sub: subiculum.

**B)** Epifluorescence images showing tdTomato expression in two mice that were injected with diluted AAV-CKII-Cre *in utero* at E13 followed by a large AAV1-CAG-FLEX-tdTomato injection in the adult hippocampus with a microliter syringe (#75, Hamilton Company). Arrows indicate parts of the injection track (i.e. not necessarily where the needle was located during the injection itself). Images were acquired as z-stacks and the Extended Focus Module in the Zeiss Zen Blue imaging software used to create a flat image. DG: dentate gyrus.

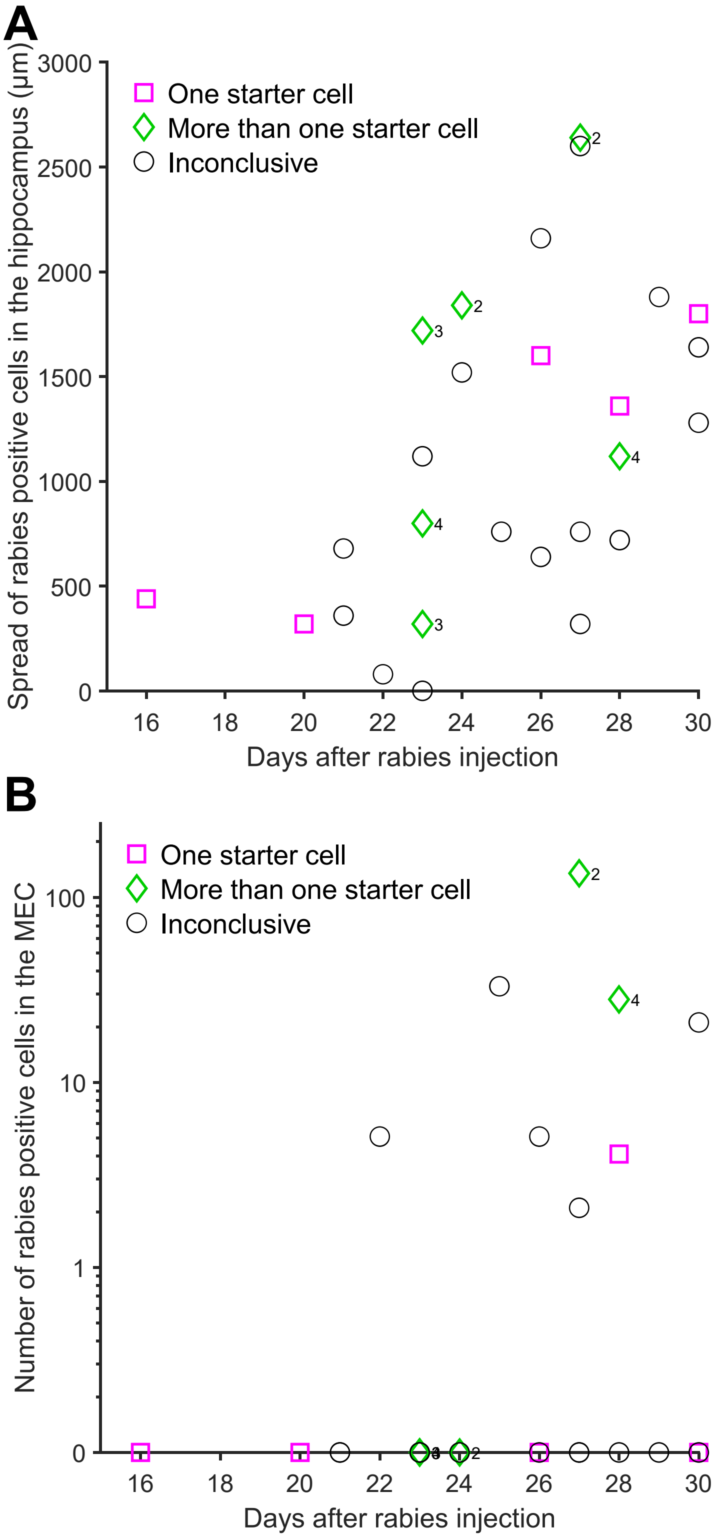


**Figure S2: Spread of rabies-positive cells in the hippocampus and number of rabies-positive cells in the MEC, related to figure 2**

**A)** The anatomical spread of rabies-positive cells along the longitudinal axis in the dorsal hippocampus increases as a function of days after the rabies injection (r = 0.54, p = 0.0036, Spearman's rho, n = 27 animals). Spread is calculated based on the number of sections in which rabies-positive cells were observed but does not include the ventral hippocampus (i.e. the spread is likely higher for animals with a long survival time). Animals with zero starter cells are not included as rabies-positive cells are absent.

**B)** The total number of rabies-positive cells observed in the ipsilateral medial entorhinal cortex (MEC) as a function of days after the rabies injection. Numbers next to diamonds indicate the number of starter cells in the 'More than one starter cell' category. Note: the y-axis follows a log scale and animals with zero starter cells are not included as rabies-positive cells are absent.

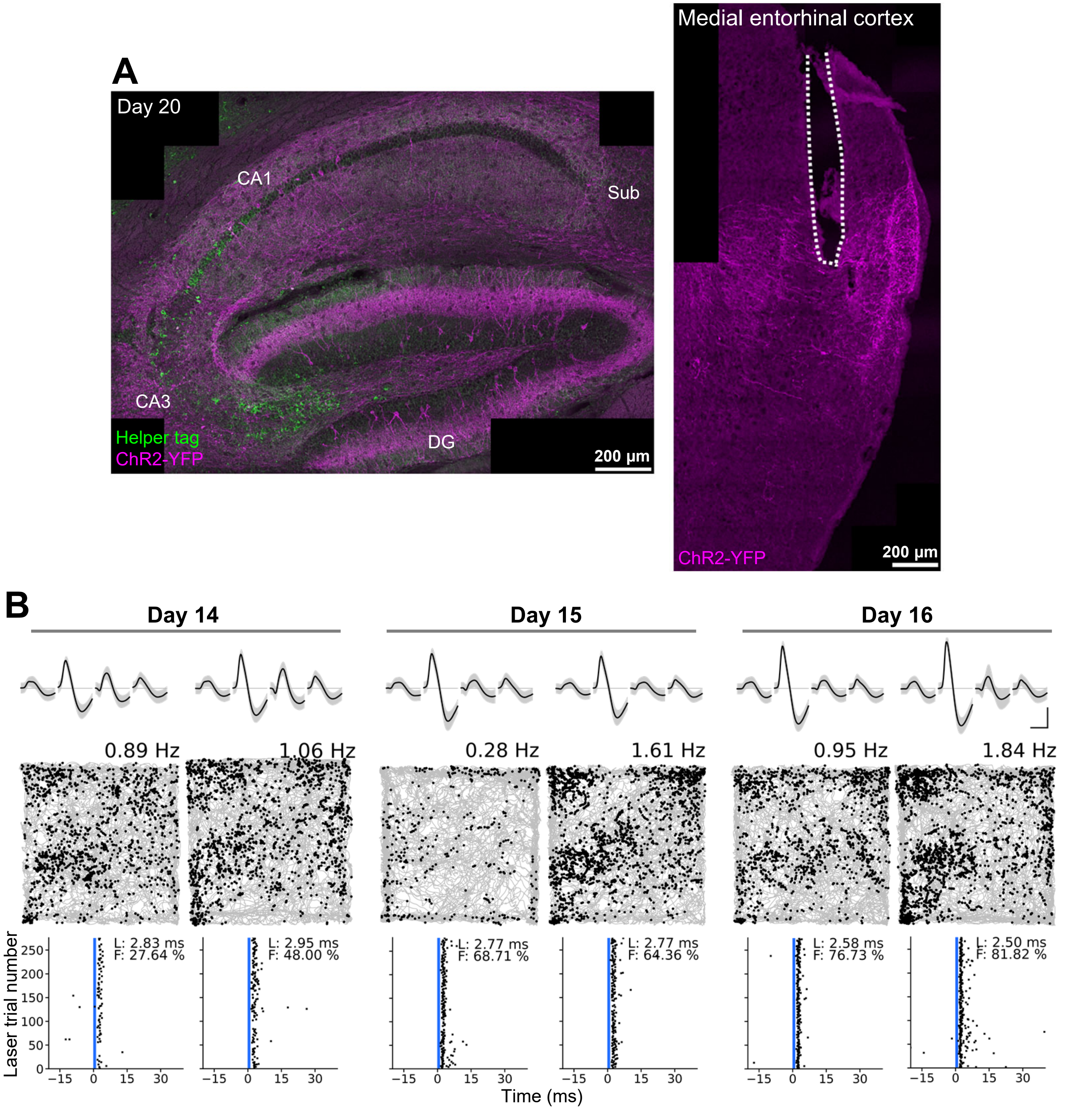


**Figure S3: Functional properties of an input cell over time, related to figure 5**

**A)** A large population of starter cells in the hippocampus was targeted by mixing the Cre virus and the helper virus before injection in the adult hippocampus. In a separate, subsequent surgery, a rabies virus carrying ChR2-YFP was injected into the hippocampus, and tetrodes with an optic fibre attached ('optrode') implanted by the ipsilateral MEC (as in **Figure 5**). Left: confocal maximum intensity projection of the hippocampus showing cells expressing helper proteins and/or ChR2-YFP 20 days after the rabies injection. Right: confocal maximum intensity projection showing part of the tetrode track (indicated by white dotted line) and expression of ChR2-YFP in the MEC.

**B)** Waveforms (top) and path plots (middle) of a cell that was recorded in six different sessions across three days (14, 15, and 16 days after the rabies injection). Subsequent laser trials (1 ms pulses at 10 Hz and 7.5 mW, indicated by vertical blue line) in a separate holding box (as in **Figure 5**) showed fast, robust and reliable responses to laser stimulation (bottom), indicating that this cell expresses ChR2 and provides monosynaptic input to the hippocampus. Grey lines indicate the path of the mouse in the open field while black dots indicate locations (path plots) or time relative to laser stimulation (laser stimulation plots) at which units from the cell was recorded. Scale bars for waveform plots: 0.5 ms (horizontal) and 50 μV (vertical). Values above path plots show the average firing rate in the open field. L: latency, F: fidelity.

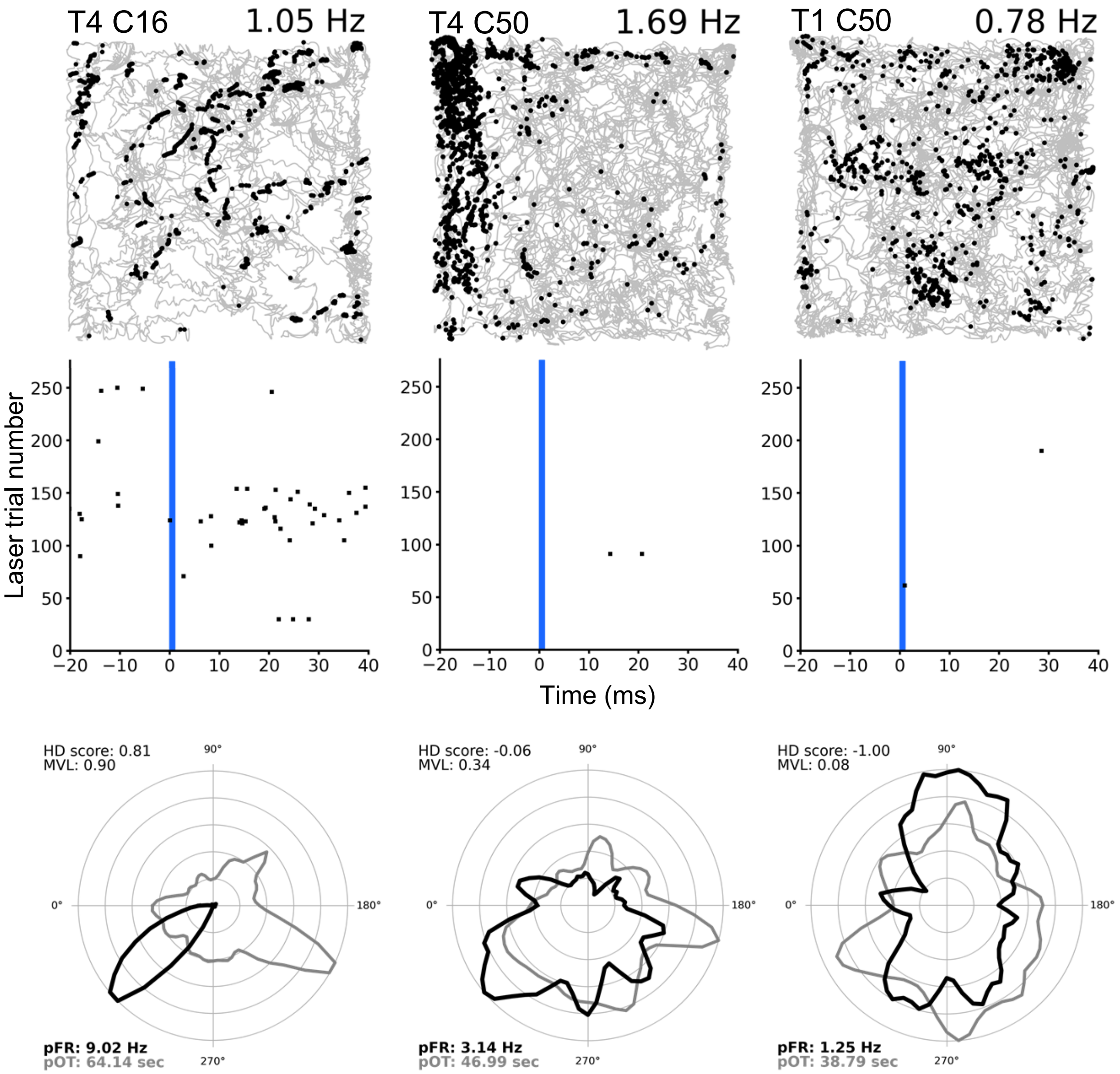


**Figure S4: Non-responsive cells, related to figure 5**
Path plots (top) of three different example cells recorded in the same area as those depicted in **Figure 5**, but that do not respond to laser stimulation (middle, blue vertical line). Head direction (HD) plots from the open field recordings are also shown (bottom, firing rate (black) and occupancy time (grey) are normalised to their own maximum in each session). The cells shown include a head direction cell (left, 17 days after rabies injection), a border cell (middle, 18 days after rabies injection) and a grid cell (right, 19 days after rabies injection). Grey lines in the open field indicate the path of the mouse while black dots indicate locations (path plots) or time relative to laser stimulation (laser stimulation plots) at which units from an individual cell was recorded. Right-centered values above path plots show the average firing rate in the open field. 1 ms laser pulses were delivered at 10 Hz and with 7.5 mW power. T: tetrode number, C: cell ID, MVL: mean vector length, pFR: peak firing rate, pOT: peak occupancy time.
