## Supplemental tables with legends for "All-viral tracing of monosynaptic inputs to single birthdate-defined neurons in the intact brain"

| Animal ID | Sex | Cre dilution | Helper virus (nl) | Days with rabies | TSA+ used? | Starter cells | Starter cell identity | Rabies cells in hipp | Rabies cells in EC | Figure |
|---|---|---|---|---|---|---|---|---|---|---|
| **59457** | **M** | **1000** | **41.4** | **16** | **no** | **1** | **CA3** | **8** | **0** | **2, 3, S2** |
| **60047** | **F** | **None** | **82.8** | **20** | **no** | **1** | **GC** | **2** | **0** | **2, 4, S2** |
| 61092 | F | 500 | 55.2 | 21 | yes | Inconclusive | | 2 | 0 | 2, S2 |
| 61144 | M | 500 | 55.2 | 21 | yes | Inconclusive | | 4 | 0 | 2, S2 |
| 61159 | M | 500 | 55.2 | 22 | yes | 0 | | 0 | 0 | 2, S2 |
| 61164 | M | 500 | 55.2 | 22 | yes | Inconclusive | | 3 | 5 | 2, S2 |
| 59930 | F | 500 | 147.2 | 23 | no | 3 | GC, CA, sl | 12 | 0 | 2, S2 |
| 59931 | F | 500 | 101.2 | 23 | no | 4 | 2xGC, CA1, Sub | 38 | 0 | 2, S2 |
| 59944 | F | 500 | 46 | 23 | no | 3 | 2xCA1, CA1sr | 11 | 0 | 2, S2 |
| 59946 | F | 500 | 92 | 23 | no | Inconclusive | | 30 | 0 | 2, S2 |
| 59975 | M | 500 | 115 | 23 | no | Inconclusive | | 1 | 0 | 2, S2 |
| 60650 | F | 500 | 69 | 24 | yes | 2 | CA, WM | 101 | 0 | 2, S2 |
| 60651 | F | 500 | 55.2 | 24 | yes | Inconclusive | | 33 | 0 | 2, S2 |
| 59943 | M | 500 | 46 | 25 | no | 0 | | 0 | 0 | 2, S2 |
| 59977 | M | 500 | 55.2 | 25 | no | Inconclusive | | 32 | 33 | 2, S2 |
| 59979 | F | 500 | 55.2 | 26 | no | Inconclusive | | 7 | 0 | 2, S2 |
| **60652** | **F** | **500** | **55.2** | **26** | **yes** | **1** | **GC** | **36** | **0** | **2, 4, S2** |
| 60821 | M | 500 | 55.2 | 26 | yes | Inconclusive | | 116 | 5 | 2, S2 |
| 60222 | F | 500 | 55.2 | 27 | no | Inconclusive | | 72 | 0 | 2, S2 |
| 60221 | F | 500 | 55.2 | 27 | no | Inconclusive | | 50 | 2 | 2, S2 |
| 60822 | M | 500 | 55.2 | 27 | yes | Inconclusive | | 7 | 0 | 2, S2 |
| 60823 | F | 500 | 55.2 | 27 | yes | 2 | CA3sr, CA3 | 668 | 135 | 2, S2 |
| **60259** | **F** | **500** | **55.2** | **28** | **yes** | **1** | **DG** | **35** | **4** | **2, 4, S2** |
| 60824 | F | 500 | 55.2 | 28 | yes | 4 | GC, CA3, WM, CA1so | 74 | 28 | 2, S2 |
| 60825 | F | 500 | 55.2 | 28 | yes | Inconclusive | | 54 | 0 | 2, S2 |
| 60218 | M | 500 | 55.2 | 29 | no | 0 | | 0 | 0 | 2, S2 |
| 60919 | F | 500 | 55.2 | 29 | yes | Inconclusive | | 8 | 0 | 2, S2 |
| 60920 | F | 500 | 55.2 | 29 | yes | 0 | | 0 | 0 | 2, S2 |
| **60219** | **M** | **500** | **55.2** | **30** | **no** | **1** | **CA so** | **27** | **0** | **2, 3, S2** |
| 60389 | M | 500 | 55.2 | 30 | no | Inconclusive | | 74 | 21 | 2, S2 |
| 60390 | M | 500 | 69 | 30 | yes | Inconclusive | | 97 | 0 | 2, S2 |

**Table S1: Experimental details and analysis of animals used for single cell experiments**
All animals were injected with Cre at E13 and animals with a single starter cell are highlighted in bold. The number of rabies positive cells in the dorsal hippocampus include starter cells. The table is sorted by the ‘Days with rabies’ column. All CA/CA3/CA2/CA1 cells are in the pyramidal cell layer unless indicated to be in so or sr.
hipp: dorsal hippocampus, EC: entorhinal cortex, CA: cell in theCA3/CA2/CA1 border area, GC: granule cell, Sub: cell in the subiculum, WM: cell in the white matter, sl: cell in stratum lucidum, so: cell in stratum oriens, sr: cell in stratum radiatum

| Animal ID | Sex | Embryonic day for Cre injection | Cre dilution | Helper virus (nl) | Days with rabies | Experiment | Figure |
|---|---|---|---|---|---|---|---|
| 72359 | M | N/A | N/A | N/A | 14 | WT with rabies | S1 |
| 58695 | M | E12 | None | 404.8 | 14 | Cre, N2cG and rabies | S1 |
| 58482 | M | E13 | 500 | N/A | N/A | Check of Cre expression | S1 |
| 58483 | M | E13 | 500 | N/A | N/A | Check of Cre expression | S1 |
| 70375 | M | N/A (adult injection) | N/A | 151.8 | 20 | Optogenetics | S3 |
| 58313 | F | E12 | None | 404.8 | 30 | Optogenetics | 5, S4 |
| 58253 | F | E14 | None | 404.8 | 24 | $Ca^{2+}$ imaging | 6 |
| 60388 | M | E13 | 500 | 55.2 | 34 | $Ca^{2+}$ imaging | 7 |

**Table S2: Experimental details of animals used for control and functional experiments**
WT: wildtype (i.e. mouse not injected with Cre *in utero*). The TSA Plus staining protocol was not used with any of these animals.
